# Disruption of Nrxn1α within excitatory forebrain circuits drives value-based dysfunction

**DOI:** 10.1101/818419

**Authors:** Opeyemi Alabi, Mara Robinson, Michael Fortunato, Joe W. Kable, Marc V. Fuccillo

## Abstract

Goal-directed behaviors, complex action sequences that maximize reward, are essential for normal function and are significantly impaired across neuropsychiatric disorders. Despite extensive associations between genetic mutations and these brain disorders, the mechanisms by which candidate genes contribute to goal-directed dysfunction remains unclear, owing to challenges in (1) describing aspects of reward processing that drive goal-directed dysfunction, (2) localizing these deficits to specific brain circuits and (3) relating changes in physiology to behavioral alterations. Here we examined mice with mutations in Neurexin1α, a presynaptically-localized adhesion molecule with widespread neuropsychiatric disease association, in value-based decision-making paradigms. We found that Neurexin1α knockout animals exhibited blunted choice bias towards outcomes associated with greater benefits. Mutant mice were similarly impaired in avoiding costlier, benefit-neutral actions. Analysis of trial-by-trial choice data via reinforcement learning models suggested these behavioral patterns were driven largely by deficits in the updating and representation of choice values. Employing conditional gene ablation and region-specific Cre-recombinase strains, we revealed that Neurexin1α disruption within forebrain excitatory projection neurons, but not thalamic population s, recapitulated most aspects of the whole-brain knockout phenotype. Finally, utilizing *in vivo* recordings of direct pathway spiny neuron population calcium activity, we demonstrated that selective knockout of Neurexin1α within forebrain excitatory neurons disrupts reward-associated neural signals within striatum, a major site of feedback-based learning. By relating deficits in value-based decision-making to region-specific Nrxn1α disruption and changes in reward-associated neural activity, we reveal potential neural substrates for the pathophysiology of neuropsychiatric disease-associated cognitive dysfunction.

## INTRODUCTION

The ability to construct and perform action sequences oriented towards a desired goal is a critical aspect of animal fitness. The execution of goal-directed behaviors engages numerous cognitive and motor processes – reward-relevant cues from the external world must be accurately represented, internal motivational states must be weighed and these diverse input variables integrated by circuits that control eventual motor output (Doya, 2008). Cortical brain regions including the orbital frontal cortex (OFC), medial prefrontal cortex (mPFC) and anterior cingulate cortex (ACC) represent aspects of reward value and history (Bari et al., 2019; Bartra et al., 2013; Euston et al., 2012; Noonan et al., 2011; Padoa-Schioppa and Conen, 2017; Rushworth et al., 2011; Rushworth et al., 2012), primary sensory cortices and midline thalamic nuclei represent reward-associated environmental signals (Parker et al., 2019; Znamenskiy and Zador, 2013), and motor thalamic nuclei ensure the smooth performance of actions (Diaz-Hernandez et al., 2018). Furthermore, the flexible adaptation of action-associated value signals is necessary in dynamically changing environments and is supported by error-monitoring signals within the ACC and basolateral amygdala, as well as reward prediction errors encoded by striatal-targeting midbrain dopaminergic neurons (McGuire et al., 2014; Schultz et al., 1997; Ullsperger et al., 2014; Yacubian et al., 2006). The dorsal striatum, as a major integration site for projections from these areas, has critical functions in action selection, motor performance and reinforcement learning (Balleine et al., 2007; Cox and Witten, 2019; Lee et al., 2015; Vo et al., 2014).

While neuropsychiatric disorders are phenotypically distinct, they are characterized by overlapping deficits in behavioral domains encompassing social, attentional and cognitive function (Ferrante et al., 2019). Deficits in goal-directed decision making, and specifically in how reward shapes selection of actions, are a core endophenotype shared across disorders, including schizophrenia, autism spectrum disorders (ASD), obsessive compulsive disorder and Tourette Syndrome (Barch and Dowd, 2010; Corbett et al., 2009; Dichter et al., 2012; Dowd et al., 2016; Gillan and Robbins, 2014; Griffiths et al., 2014; Hill, 2004; Maia and Frank, 2011; Solomon et al., 2015). While initial assumptions of blunted hedonic processing in schizophrenia have not been widely substantiated (Gold et al., 2008), other alterations of reward processing have been uncovered, including increased temporal decay of reward value (Ahn et al., 2011; Barch and Dowd, 2010; Heerey et al., 2007; Yu et al., 2017) and impairments in action-outcome learning (Gold et al., 2015; Morris et al., 2015). Disruptions in trial-to-trial feedback-based learning in this disorder may reflect perturbations to reinforcement learning error signals or the manner in which they are integrated to alter choice (Hernaus et al., 2019; Hernaus et al., 2018). Though typically associated with restricted patterns of interest and social reward impairments, recent studies have also revealed reinforcement learning deficits in ASD patients as well (Hill, 2004; Solomon et al., 2015). Interestingly, ASD patients revealed selective impairment of choice accuracy on high probability stimulus pairs, marked by decreases in win-stay choice patterns (Solomon et al., 2015).

As these diverse goal-directed phenotypes may arise from common underlying neural circuit mechanisms, exploring analogous reinforcement behaviors in tractable rodent models may provide substantial circuit level insights into key drivers of behavioral pathology (Griffiths et al., 2014). Given the wealth of potential genetic models arising from recent genetic association studies (De Rubeis et al., 2014; Willsey et al., 2013; Willsey and State, 2015), our approach was to focus on a gene conferring risk for multiple neuropsychiatric disorders – in effect, a molecule whose disruption broadly sensitizes the brain to subsequent, disease-specific insults. Neurexin1α (Nrxn1α is an evolutionarily conserved cell adhesion molecule, in which rare *de novo* mutations and single nucleotide variations confer significant risk for ASDs, schizophrenia, Tourette Syndrome and Obsessive-Compulsive Disorder (Ching et al., 2010; Duong et al., 2012; Huang et al., 2017; Kirov et al., 2009; Rujescu et al., 2009). Nrxn1α functions as a presynaptic hub for transynaptic binding of numerous postsynaptic functions as a ners at both part excitatory and inhibitory synapses (Missler et al., 2003; Sudhof, 2017). These interactions are thought to mediate the initial specification and long-term integrity of synapses (Anderson et al., 2015; Aoto et al., 2013; Chubykin et al., 2007; Krueger et al., 2012; Soler-Llavina et al., 2011; Sudhof, 2017; Varoqueaux et al., 2006). Nrxn1α is broadly expressed throughout the brain, but particularly enriched in cortico-stri al-thalamic loops proposed to govern motor control, action selection and reinforcement learning (Fuccillo et al., 2015; Ullrich et al., 1995).

Documented behavioral abnormalities in Nrxn1α knockout animals are numerous, including reduced nest building and social memory, increased aggression and grooming, enhanced rotarod learning, and reductions in pre-pulse inhibition (Dachtler et al., 2015; Esclassan et al., 2015; Etherton et al., 2009; Grayton et al., 2013). Furthermore, work from Nrxn1α knockout rats described male-specific reductions in operant responding under increasing variable interval responding schedules (Esclassan et al., 2015). While these studies paint a broad picture of behavioral dysfunction, it has been challenging to capitalize on these observations to reveal Nrxn1α ’s underlying mechanistic contribution to disease-relevant behavioral outcomes. Thi s hurdle may reflect our poor understanding of the specific computational algorithms and neural circuit implementations for the behavioral states interrogated by the aforementioned behavioral assays.

In this paper, we uncover widespread alterations in reward processing in mice with Nrxn1α mutations, manifest both as inefficient choice patterns and blunted modulation of task engagement. These deficits were observed consistently across a range of value comparisons and feedback rates, suggesting trait-like characteristics of mice with Nrxn1α disruption. Modeling of choice patterns suggests that altered choice behavior relates to a specific neurocognitive domain – the learning and representation of choice values in the face of new reward information. To uncover causal circuits for this reward processing defect, we performed cell type-specific ablations of Nrxn1α. We found that the brain-wide mutant behavioral deficits can be largely recapitulated with Nrxn1α disruption in excitatory forebrain projection neurons, but not thalamic nuclei. Finally, we demonstrate dysregulation of value-associated circuit dynamics in direct pathway neurons of the dorsomedial striatum accompanying forebrain projection neuron-specific Nrxn1α disruption. This work represents an important step in characterizing the genetic contributions to circuit dysfunction for a core neuropsychiatric disease-relevant behavior – how animals shape their actions according to cost and benefit.

## RESULTS

### Neurexin1α mutants acquire a simple goal-directed task

To study value-based action selection in Nrxn1α mutant mice, it was important to establish whether these mice exhibit baseline deficits in acquisition and performance of simple instrumental tasks. A previous study uncovered instrumental learning deficits in Nrxn1α knockout (KO) rats, characterized by an inability to increase responding levels under reward interval uncertainty (Esclassan et al., 2015). To explore this, we ran behaviorally naïve Nrxn1α wildtype and littermate KO animals in a self-initiated, light-cued choice task (Fig.S1 A). Over 10 sessions, mice learned to initiate trials via lit central nosepoke, triggering a 10-second choice period in which one of two lateralized pokes was illuminated. Selection of the lit port resulted in reward, while pokes into the other two unlit ports resulted in time out. We assessed performance by measuring the proportion of choices allocated towards the reward port and estimated the willingness of animals to work for reward as the total number of center port trial initiations within a one-hour session. Nrxn1α mice exhibited robust task engagement (Fig.S1B) and accuracy curves that were statistically indistinguishable from wildtype controls (Fig.S1C). Thus, Nrxn1α loss-of-function does not lead to broad impairments in the execution of simple goal -directed responding under low schedule requirements.

### Neurexin1α KOs have blunted responses to differential reward outcomes

To specifically test how Nrxn1α mutant mice use value information to guide future choice, we employed a feedback-based paradigm that samples mouse choice across outcome values (Fig.1A). This task utilizes a block design with dynamically alternating reward contingencies triggered by recent choice biases to maintain outcome sensitivity over hundreds of trials (Fig.1A, see (Alabi et al., 2019) and STAR Methods for details). Mice self-initiated trials in which two lateralized (left/right) choices with contrasting reward volumes were available. To broadly cover value space, we tested 4 relative reward ratios, in conditions with both high (P_rew_=0.75) and low (P_rew_=0.4) feedback rates. Global performance in this task was significantly altered by the relative magnitude of rewarded outcomes for both wildtype and KO animals, demonstrating that larger reward contrasts drive more biased choice patterns (Fig.1B). We observed a significant decrement in session performance across relative reward contrasts in Nrxn1α KO mice as compared to wildtype (Fig.1B).

**Figure 1.**
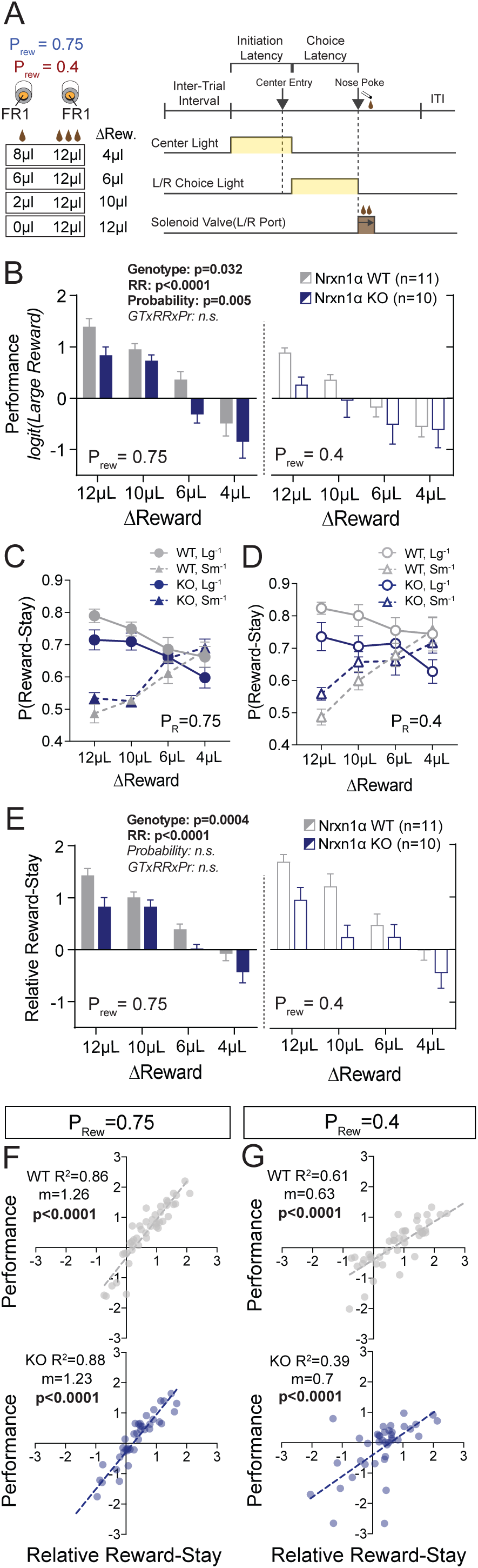
Neurexin1α disruption leads to deficits in value-based selection of actions. (A) Schematic of trial structure wherein mice perform repeated self-initiated trials with contrasting reward volumes associated with each port. Animals were tested at four relative reward ratios across high (P_rew_=0.75) and low (P_rew_=0.4) reinforcement rates. See STAR Methods for details. (B) Both probability of reinforcement and volume difference modulate the probability at which mice select the large reward option. Nrxn1α KOs (blue, n=10) select the high benefit alternative at a lower rate than their WT littermates (gray, n=11) across reward environments (3-way RM-ANOVA). (C,D) For both WT and KO animals, the relative magnitude of rewarded outcome has a significant effect on the stay-probability for that alternative. (E) The relative reward stay (RRS), which quantifies the relative tendency of animals to repeat choices after specific outcomes, was sensitive to relative magnitude of rewards but not reward probability. In comparison to WT littermates, Nrxn1α KOs less dynamically alter their choice behavior after large reward outcomes than small reward outcomes (3-way RM-ANOVA). (F,G) The RRS is a significant predictor of session performance for both WT and KO mice at both rates of reinforcement. Note RRS is a better predictor of task performance at high reinforcement rates, reflecting the preponderance of unrewarded outcomes in low reinforcement conditions. All data represented as mean ± SEM.

Performance in this task can be altered by changes in: (1) the integration window for feedback history; (2) sensitivity to outcome feedback; (3) response to the absence of feedback; (4) flexibility to changing contingencies (Alabi et al., 2019). To assess whether Nrxn1α KOs show altered influence of reward history on current choice, we employed logistic regression models to generate weighted coefficients representing the relative effect of choice and outcome 5 trials preceding current choice (Lau and Glimcher, 2005; Parker et al., 2016; Tai et al., 2012). Similar to prior work (Alabi et al., 2019), we found that wildtype mice and Nrxn1α KOs heavily discount all but the immediately preceding trial (t-1) (Fig.S1E-H). This suggested a significant portion of variability in choice behavior could be understood by analyzing the results of the t-1 trial.

To investigate whether the observed Nrxn1α KO deficits relate to alterations in feedback response, we analyzed patterns of choice following specific outcomes. We reasoned the efficiency of selecting more beneficial alternatives is driven by comparative sensitivity to reward outcomes of differing magnitudes. Given the steep decay of our logistic regression curves, we calculated the relative reward-stay (RRS), a descriptive measure of the relative reinforcing properties of large and small rewarded t-1 outcomes (previously called relative action value in (Alabi et al., 2019)). We noted smaller gaps between large reward-stay and small reward-stay behavior in Nrxn1α KOs as compared with wildtypes (Fig.1C,D), leading to smaller relative reward-stay values across reward contrasts and feedback environment (Fig.1E). The significant correlation between RRS and performance across genotypes highlights the importance of relative reward reinforcement on global performance (Fig.1F,G). In sum, constitutive Nrxn1α KO mice exhibit a reduced ability to bias their choices in response to differing reward feedback magnitudes, and this deficit in part underlies the observed differences in global behavioral efficiency.

### Neurexin1α mutants demonstrate normal levels of cognitive flexibility

Deficits in behavioral adaptability have been observed across neuropsychiatric disorders and can impact performance in our task (Alabi et al., 2019). To explore differences in flexible responding between Nrxn1α mutants and wildtypes, we compared choice patterns immediately before and after un-c ued contingency switches triggered by a fixed directional bias (8/10 selections to the large reward choice). This analysis revealed a strong trend suggesting Nrxn1α mutants less dynamically alter their choices in dynamic environments (Fig.S1I). To assess whether the observed differences in adaptability are derived from deficits in cognitive flexibility or secondary to alterations in value-based choice behavior, we utilized extra-dimensional set shifting and ego-centric spatial reversal tasks. Given that performance of these tasks is thought to engage distinct cortico-striatal circuits (Floresco et al., 2008; Ghods-Sharifi et al., 2008), we felt testing animals in both would thoroughly interrogate potential cognitive flexibility deficits. Interestingly, we observed no deficit in the ability of Nrxn1α KO mice to adapt to different rule structures (Fig.S1J) or their ability to modulate their responding to alterations in existing rule structures (Fig.S1K). This suggests that adaptability deficits observed in our value-based serial reversal task are not caused by global cognitive inflexibility, but may be secondary to deficits in outcome sensitivity.

### Neurexin1α mutants exhibit abnormalities in outcome-related task engagement

The temporal relationship between action and reinforcement modulates the degree to which rewards shape behavior. To assess whether observed differences in outcome sensitivity resulted from divergent temporal patterns of task performance, we analyzed relevant task latencies. In our experiments, we noted that the latency to initiate trials increased as the cumulative volume of reward received during a given session increased (Fig.S2A). As such, we used this value as a proxy of task engagement – a reflection of the animals’ motivation or attention towards the current task (Bari et al., 2019; Hamid et al., 2016). We observed no significant discrepancies in this measure between Nrxn1α wildtype and KO mice across varied reward environments (Fig.2A). This suggests that the blunted outcome-associated choice response is not attributable to global disengagement with task events.

**Figure 2.**
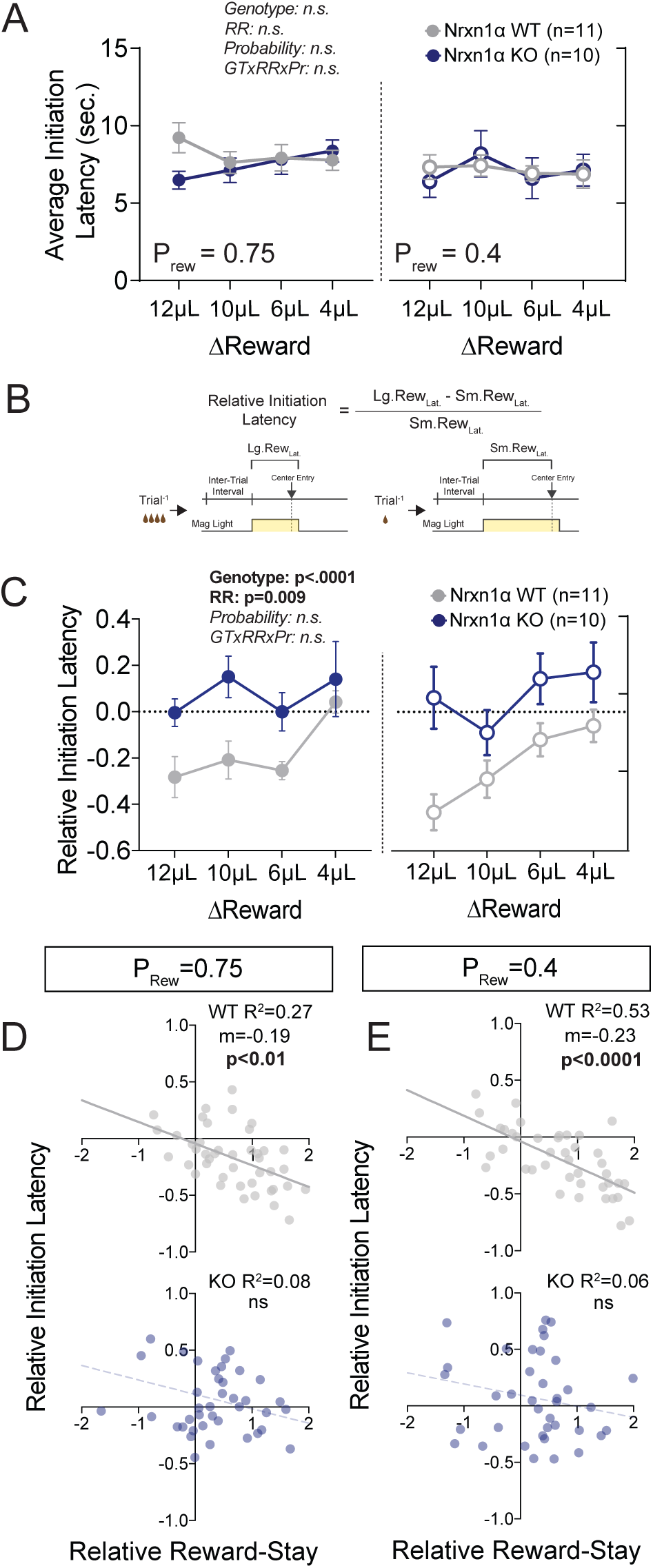
Neurexin1α mutants display altered outcome-dependent task engagement. (A) Task engagement was measured as the average latency from trial onset (center-light ON) to initiation. Nrxn1α KOs (blue, n=10) do not exhibit global deficits in task engagement in comparison to WT animals (gray, n=11) (3-way RM ANOVA). (B) Relative latency to initiate is a standardized comparison of initiation latencies following large rewarded outcomes and small rewarded outcomes within individual animals. (C) Nrxn1α WT mice modulate their trial-by-trial engagement in response to different rewarded outcomes, initiating trials more quickly after large reward outcomes than small reward outcomes. Nrxn1α KOs don’t exhibit this outcome-sensitive modulation of task engagement (3-way RM ANOVA). (D,E, top) There is a significant relationship between the ability of WT mice to select actions in response to reward discrepancy (RRS) and their ability to upregulate task engagement (relative initiation latency) which is lost in KOs (D,E bottom). All data represented as mean ± SEM.

Recent evidence suggests choice value can also modulate the vigor with which selected actions are performed (Bari et al., 2019; Hamid et al., 2016). If inefficient choice patterns of Nrxn1α observe changes KOs result from disrupted value encoding, one might expect to also observe changes in how local reward environment modulates vigor of action performance. To explore this, we compared outcome-dependent initiation latencies after large reward and small reward outcomes (Fig.2B). Interestingly, the relative latency to initiate trials in wildtype animals was significantly modulated by the relative reward ratio (Fig.2C), with animals initiating trials more quickly after large reward outcomes than small reward outcomes. In wildtype mice, larger reward contrasts produced more pronounced modulation of initiation latencies (Fig.2C, grey). In contrast, Nrxn1α knockout mice were entirely unable to modulate initiation latency in response to the magnitude of previous reward (Fig.2C, blue). The strong inverse correlation between relative reward-stay and initiation latency was lost in Nrxn1α KO mice (Fig.2D,E). Thus, while there is no difference in average task latencies between wildtypes and mutants, Nrxn1α loss-of-function disrupts the manner in which outcomes modulate local task engagement.

In addition to differences in reward-related modulation of initiation latencies, we noted a significant increase in choice latency in Nrxn1α mutants across reward environments Fig.S2B). While several studies suggest that choice latencies are correlated with choice uncertainty (Gage et al., 2010; Redish, 2016), further analysis did not reveal a reward-related component to this behavioral phenotype (not shown). This phenotype was consistently observed for each Nrxn1α loss-of-function manipulation examined here and stands in contrast to the hyperactivity previously described (Etherton et al., 2009).

### Value processing abnormalities in the Neurexin1α mouse extend to cost-based decision making

Because unrewarded trials are evenly distributed between large and small reward choices, lose-switch behavior does not influence overall task efficiency. However, we found that Nrxn1α mutant animals perform fewer choice switches after random reward omission than their wildtype littermates (Fig.S3A), suggesting Nrxn1α KOs might have deficits in using negative outcomes to influence behavior. To explore t his, we introduced mice to an alternate paradigm that modulated the value of choices using differential costs. Given the existing deficits in processing value differences probed by volume, each choice alternative was assigned a differential repetitive motor requirement (FR3 vs. FR1), but equal reward volume (Fig.3A). Reward contingencies in this paradigm were not alternated and mice were challenged to allocate their choices to the lower cost (i.e., higher value) alternative over 150 trials. After 75 trials of reward feedback, mice achieved a steady-state response pattern, with no significant changes in selection of the low-cost alternative. Interestingly, Nrxn1α KO mice do not select low-cost alternatives as frequently as wildtype littermates, bo th during sampling and steady-state periods (Fig.3B). While we noted the KOs slowed more over the session (Fig.3B), no statistically significant difference in steady-state task engagement was seen. (Fig.3C). We continued to observe an effect of genotype on choice latency (Fig.3D).

**Figure 3.**
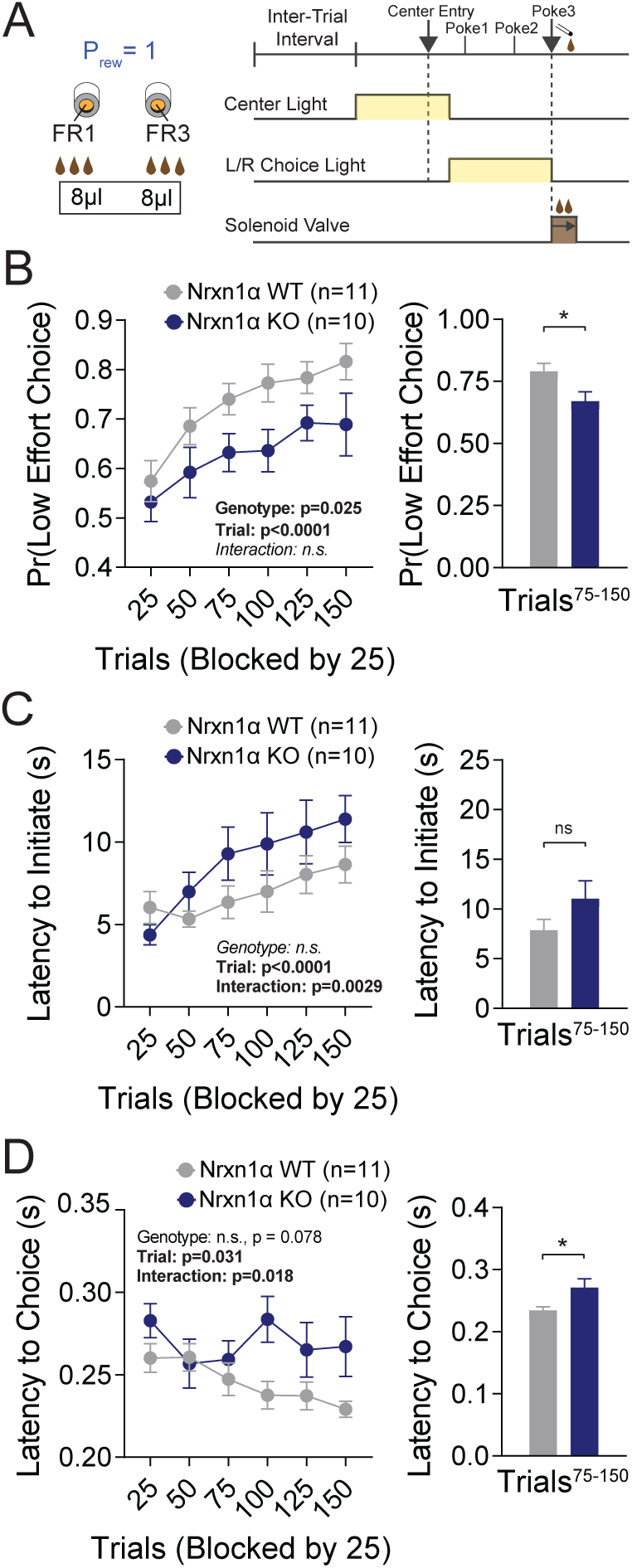
Neurexin1α mutants display a deficit in the selection of actions based on costs. (A) Effort paradigm schematic. Mice distribute choices in a session with static contingency lasting 150 trials. Animals were given choices with equal reward outcomes, but different effort requirements (FR3 v FR1). (B) Nrxn1α KOs (blue, n=10) choose less costly alternatives at a lower rate than their WT littermates (gray, n=11) (2-way RM ANOVA). The distribution of choice in both WT and KO mice is altered over the course of the block as mice acquire information about the reward contingency, with a stable difference observed over the final 75 trials (two-sample t-test *p=0.023). (C) Nrxn1α KOs exhibited a clear interaction between trial and latency to initiate, slowing as the y performed more high effort trials (2-way RM ANOVA). Nevertheless, there was no statistically significant difference in engagement at steady-state (two-sample t-test p=0.14). (D) The longer choice latencies previously described in Nrxn1α KOs was observed in steady-state responding (2-way RM ANOVA; two-sample t-tes t *p=0.017). All data represented as mean ± SEM.

### Reinforcement modeling reveals genotype-specific deficits in updating of outcome value

In order to uncover core aspects of the decision-making process underlying outcome-insensitive choice behavior in Nrxn1α mutants, we modeled action selection as a probabilistic choice between two alternatives with continually updating values (Fig.4A). We employed a modified Q-learning model with softmax decision function, including five parameters: 1) the learning rate (α), which determines the extent to which new information about state-action pairing alters subsequent behavior; 2) a reward compression parameter (γ) that captures the subjective benefit of a given reward volume; 3) a regression weight (β, the inverse temperature), linking the values of each option to the choice output; 4) a perseveration parameter (κ), capturing the effect of previous choices on subsequent choice and 5) constant terms to capture spatial biases in choice behavior (see methods for model details) (Doya, 2007; Niv, 2009; Vo et al., 2014).

**Figure 4.**
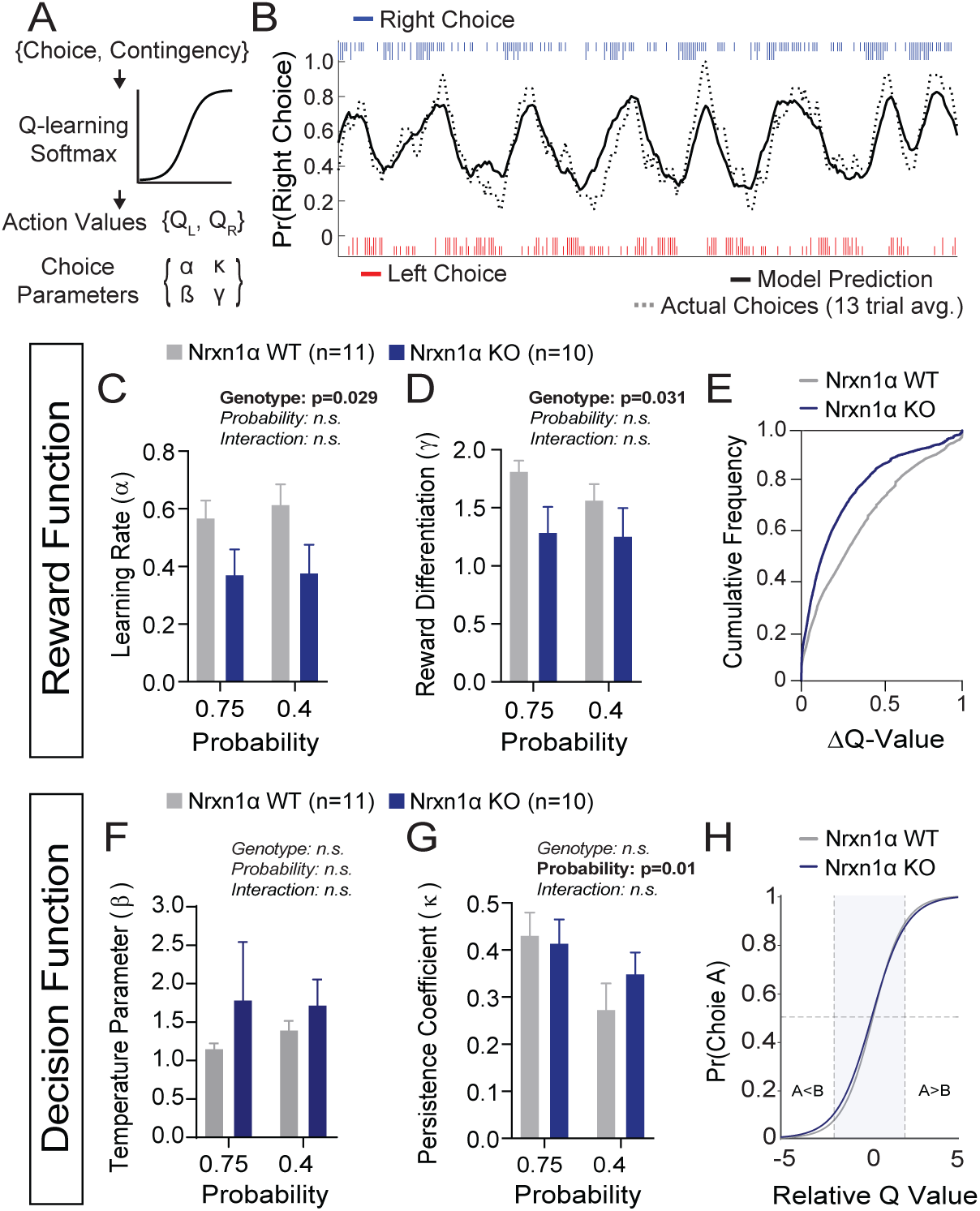
A deficit in value updating underlies abnormal allocation of choices in Neurexin1α mutants. (A) Q-learning reinforcement model. Mouse choice was modeled as a probabilistic choice between two options of different value (Q_L_,Q_R_) using a softmax decision function. Data from each reinforcement rate were grouped before model fitting. (B) Example of model prediction versus actual animal choice. Choice probability calculated in moving window of 13 trials. Long and short markers indicate large and small reward outcomes. (C,D) As compared to littermate controls (gray, n=11), Nrxn1α mutants (blue, n=10) exhibit a deficit in the learning rate, which describes the weig ht given to new reward information and, a utility function th at relates how sensitively mice integrate rewards of different magnitudes (2-way RM ANOVA). (E) Nrxn1α KOs exhibit an enrichment of low Q-value trials. (F,G) Nrxn1α mutants do not exhibi t significant differences in explore-exploit behavior (F, captured by β) or in their persistence towards previously selected actions (G, captured by β). (K) There is no significant difference in the decision function of Nrxn1α wildtype and mutant animals. All data represented as mean ± SEM. Bias figures can be found in Supplementary Fig.4.

Recent papers have demonstrated that mice exhibit stable trait-like characteristics of reward processing (Alabi et al., 2019). In light of this, we grouped the choice data of individual animals across reward ratios in order to extract stable parameters for learning, reward magnitude differentiation, choice persistence and explore/exploit behavior, as revealed by the inverse temperature parameter. We fit our model using function minimization routines and found that this model provides accurate predictions of individual animal choice patterns (Fig.4B). Separately fitting the choice data for wildtype and KO mice, we demonstrated that Nrxn1α KO mice have significantly lower α and γ parameters (Fig.4C,D), suggesting a global deficit in the updating and representation of choice values guiding decisions (Fig.4E). In contrast, we did not not observe genotypic differences for the β, κ or bias parameters (Fig.4F,G, Fig.S4), suggesting no systemic differences in how the two genotypes transform value representations into actions (Fig.4H).

### Conditional Ablation of Neurexin1α Recapitulates Value-Based Abnormalities in Forebrain Projection Neurons Recapitulates Value-Based Abnormalities

A key challenge in understanding specific molecular contributions to neuropsychiatric disease lies in uncovering neural circuits that drive relevant behavioral endophenotypes. Thus far, our findings raise the possibility that a discrete cognitive deficit in value updating underlies abnormal choice patterns exhibited by Nrxn1α KOs. Multiple cortical regions, which exhibit high expression of Nrxn1α mRNA, have been implicated in the regulation of action-outcome association and encoding of subjective values for choice (Bari et al., 2019; Euston et al., 2012; Noonan et al., 2011; Padoa-Schioppa and Conen, 2017; Rushworth et al., 2011; Rushworth et al., 2012). To test whether Nrxn1α loss-of-function in these regions was responsible for the observed reward processing deficits, we crossed a Neurexin1α conditional allele (Nrxn1α ^C^), where exon 9 was surrounded by loxP sites, to the Nex-Cre transgenic line, a strain expressing Cre-recombinase in postmitotic progenitors of forebrain projection neurons (Goebbels et al., 2006) (Fig.5A,B). Isolated mRNA from cortical dissection of Nrxn1α ^C/C^; Nex^Cre/+^ revealed a 3.5x decrease of Nrxn1α mRNA transcripts spanning exon 9 as compared to Nrxn1α C/C_;_ Nex^+/+^(Fig.5C). A modest degree of nonsense-mediated decay was also noted w ith a downstream probe (Fig.5C). The inclusion of Cre-negative interneurons (∼20% of the cortical population) and forebrain glia, both of which express Nrxn1α, likely have us underestimating the extent of Cre-mediated recombination in Nrxn1α ^C/C^; Nex^Cre/+^ mice. Given the early expression of Cre from the Nex^Cre/+^line, it is likely that the Nrxn1α ^C^ allele is recombined prior to its endogenous expression (Lukacsovich et al., 2019).

**Figure 5.**
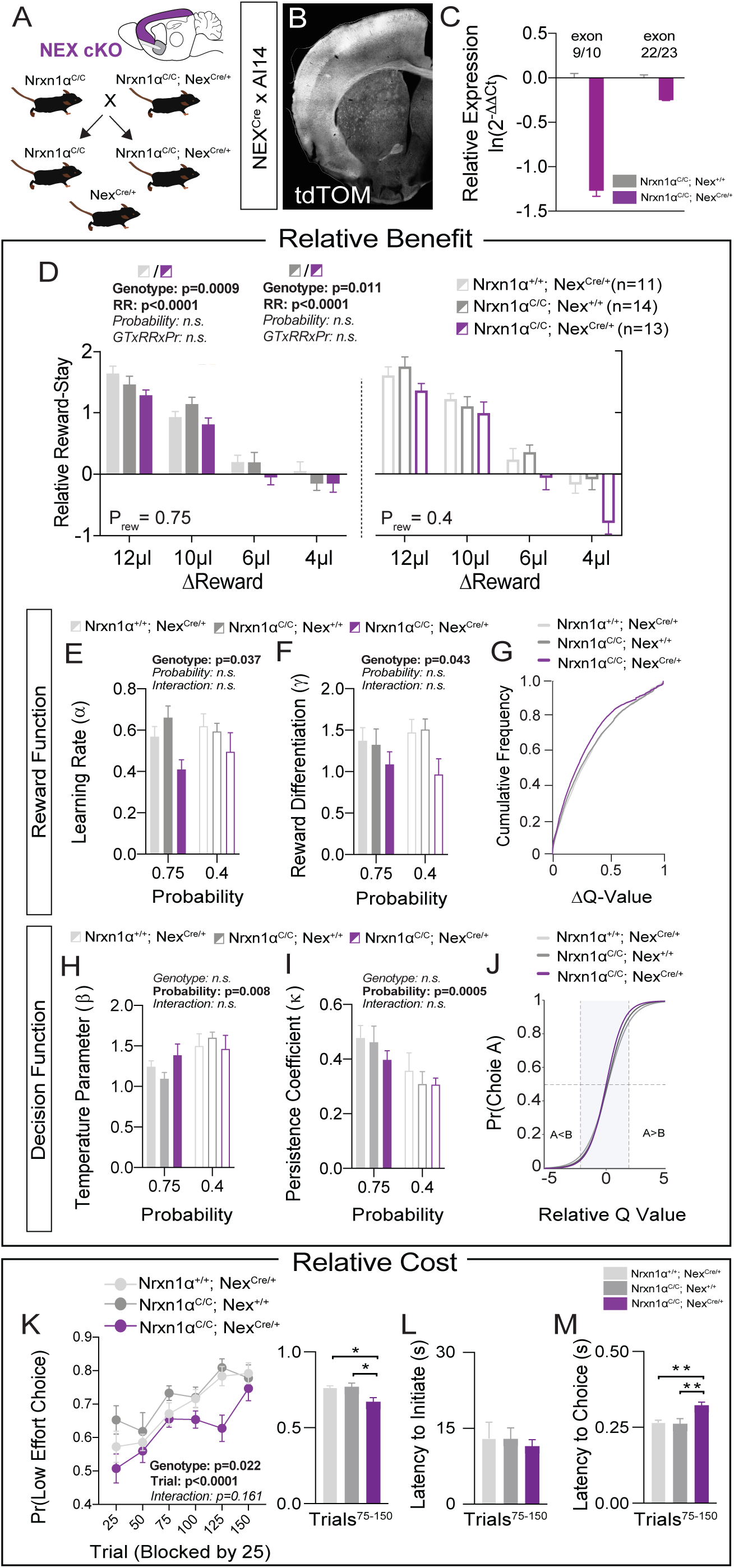
Restricted forebrain excitatory neuron deletion of Neurexin1α recapitulates choice abnormalities of constitutive KO. (A) Nrxn1α was conditionally inactivated in forebrain excitatory neurons by crossing a Nrxn1α-conditional knockout allele onto the Nex^Cre^ line. Controls for both the Nex (light gray) and Neurexin1α-conditional (dark gray) allele were analyzed. (B) Coronal section of brain from NEX^Cre/+^;Ai14 (LSL-tdTOM) reporter cross showing restriction of tdTOM fluorescence to forebrain excitatory neurons. (C) RT-qPCR of RNA from adult mouse cortex (n=3 for Nrxn1α ^C/C^;Nex^+/+^ (dark gray) and Nrxn1α ^C/C^;Nex^Cre/+^ (purple)). Cre-mediated recombination results in reduced expression of Nrxn1α mRNA detected by exon 9 probe (two-sample t-test: p<0.0001) and moderate nonsense-mediated decay (two-sample t-test: p<0.01 (D) Nrxn1α ^C/C^;Nex^Cre/+^ mutant animals (purple; n=13) exhibit a reduction in relative reward-stay as compared with Nrxn1α ^C/C^;Nex^+/+^(dark gray; n =14) and Nrxn1α ^+/+^;Nex^Cre/+^ (light gray; n=11) controls. No difference in choice allocation was observed between control animals (genotype: p=0.88, relative reward: ***p<0.0001, probability: p=0.26, 3-way interaction: p=0.25; 3-way RM ANOVA). (E,F) Similar to Nrxn1α constitutive knockouts, Nrxn1α ^C/C^;Nex^Cre/+^ mutant mice have a deficit in utilizing new re ward information to update an d represent choice values. The mutants exhibit a deficit in the learning rate (α) and in the reward volume sensitivity parameter (γ) (both γ analyzed by 2-way RM ANOVA). (G) This leads to an enrichment of low ΔQ-value trials in mutant mice. (H-J) Nrxn1α ^C/C^;Nex^Cre/+^ mutants do not differ from littermate controls for the relationship between choice value and decision behavior (H) and biases towards previous choice behavior (I). As a result, there is no significant difference in the decision function of control and mutant animals. (K-M) Nrxn1α ^C/C^;Nex^Cre/+^ mutants exhibit a deficit in the allocation of choices guided by relative choice costs (K, 2-way RM ANOVA, left; 1-Way ANOVA w/ Tukey’s multiple comparison, right, *p<0.05). Mutants exhibit no difference in task engagement (L, 1-Way ANOVA w/ Tukey’s multiple comparison, p>0.05) but recapitulate deficit in choice latencies (M, 1-Way ANOVA w/ Tukey’s multiple comparison, **p<0.01). All data represented as mean ± SEM.

In order to test the effects of Nrxn1α loss-of-function in forebrain projection neurons, we repeated the value-based tasks in Nrxn1α ^C/C^; Nex^Cre/+^mice. To account both for potential hypomorphic effects of the Nrxn1α conditional allele as well as effects of constitutive Cre expression in the Nex^Cre^ driver, we utilized two controls: Nrxn1α +/+_;_ Nex^Cre/+^and Nrxn1α ^C/C^; Nex^+/+^. We observed a significant effect of Nex^Cre^deletio n of Nrxn1α on the choice distributions of mice as compared to both control groups (Fig.5D). Similar to global Nrxn1α deletion, Nrxn1α ^C/C^; Nex^Cre/+^mutant animals were less able to bias their choice patterns towards more beneficial outcomes. We noted no significant difference in behavioral flexibility (Fig.S5A) or loss-avoidance (Fig.S5B) in these mice. Neither the Nrxn1α ^C/C^; Nex^+/+^ conditional control nor the Nrxn1α ^C/C^; Nex^Cre/+^ mutant animals displayed the reward-related modulation of initiation latencies observed in the Nrxn1α wildtype animals (Fig.S5C), precluding conclusions regarding local modulation of action vigor. Similar to constitutive Nrxn1α KOs, we noted an increased choice latency across varied reward environments (Fig .S5D).

To assess whether altered choice patterns in forebrain-specific Nrxn1α KOs resulted from the same core reward processing abnormalities seen in Nrxn1α constitutive KO mice, we again employed reinforcement modeling of choice and outcome data. As in whole-brain KOs, we observed a significant effect of genotype on both learning rate and reward discrimination parameters (Fig.5E,F), generating a shift in the distribution of action value contrasts in Nrxn1α ^C/C^; Nex^Cre/+^ mice (Fig.5G). In keeping with prior data, we observed no genotypic differences in value-related explore/exploit behavior, choice persistence or average bias (Fig.5H-J, Fig.S5E).

As a final test of whether Nrnx1α deletion from forebrain excitatory neurons could recapitulate the whole-brain KO phenotype, we tested conditional mice in our effort-based cost paradigm. Similar to the Nrxn1α constitutive KOs, the Nrxn1α C/C_; Nex_Cre/+ conditional mutants exhibited reduced selection of the lower-cost alternative than both groups of control animals (Fig.5K). Average task engagement was not abnormal in these animals (Fig.5L, Fig.S5F), but we again noted a persistent increase in choice latency (Fig.5M, Fig.S5G). Taken together, these data suggest that mid-embryonic deletion of Nrxn1α is sufficient to produce similar perturbations of reward processing as those observed in the whole-brain Nrxn1α KO mice.

### Deletion of Neurexin1α in Thalamic Nuclei Does Not Recapitulates Choice Deficits

Neurexin1α is also highly expressed in multiple subcortical regions involved in the selection and performance of goal-directed actions (Bradfield et al., 2013; Diaz-Hernandez et al., 2018; Fuccillo et al., 2015; Ullrich et al., 1995). In order to assess the specificity of forebrain excitatory Nrxn1α conditional KOs (cKOs) in driving reward processing abnormalities, we conditionally deleted Nrxn1α in developing thalamic nuclei via an Olig3-Cre driver line (Fig.6A-C) (Vue et al., 2009), reasoning that Nrxn1α loss-of-function in this region might similarly contribute to abnormal choice patterns observed in constitutive KOs. However, in contrast to the forebrain excitatory (cKOs), thalamic cKOs could not recapitulate the deficits in value processing observed in whole-brain Nrxn1α mutants (Fig.6D, Fig.S6A-D). There was no statistically significant genotypic difference in the ability to modulate choice distributions in response to reward (Fig.6D), nor in any parameters of the fitted reinforcement model (Fig.6E-J, Fig.S6E). Additionally, we noted no significant deficit in choice allocation when reward value was modulated by effort (Fig.6K-M, Fig.S6F). The only aspect of the constitutive KO phenotype partially recapitulated by the thalamic cKOs was the increased choice latency in the fixed contingency paradigm (Fig.S6G but see Fig.6M).

**Figure 6.**
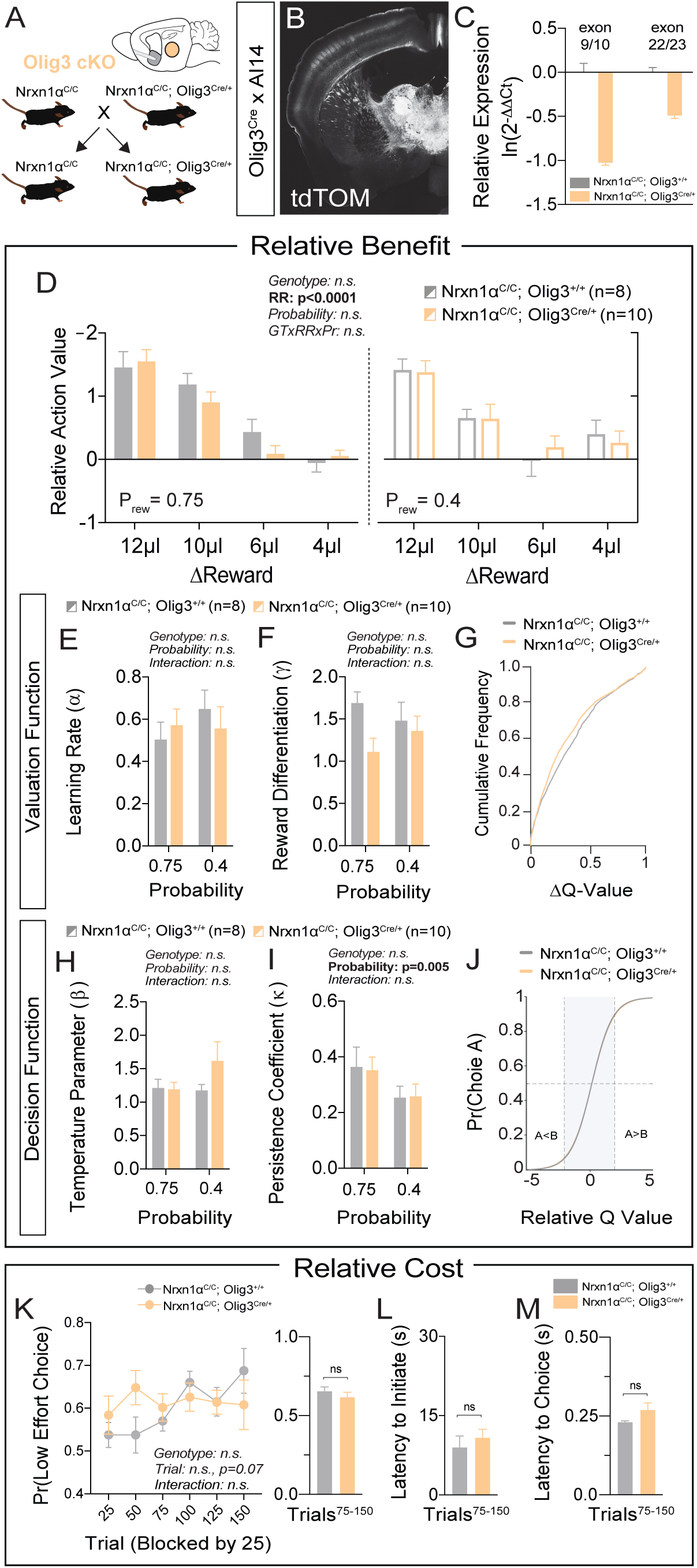
Specific deletion of Neurexin1α in thalamic nuclei does not reproduce choice deficit observed in constitutive KO. (A) Neurexin1α was conditionally inactivated in thalamic progenitor cells by crossing the Neurexin1α-conditional knockout line onto the Olig3-Cre line. (B) Coronal section of Olig3^Cre^; A i14 reporter cross showing expression of tdTOM broadly throughout thalamic nuclei. (C) RT-qPCR of RNA from adult mouse thalamus (n=2 for Nrxn1α^C/C^;Olig3^+/+^ (gray); n=3 for Nrxn1α ^C/C^;Olig3^Cre/+^(orange)). Cre-mediated recombination results in reduced expression of Nrxn1α mRNA detected by exon 9 probe (two-sample t-test: p<0.0001) and moderate nonsense-mediated decay (two-sample t-test: p<0.001) (D) Nrxn1α ^C/C^;Olig3^Cre/+^ mutant animals (orange; n=10) do not exhibit changes in relative reward-stay in comparison with Nrxn1α ^C/C^;Olig3^+/+^(gray; n=8) control animals. (E-G) Nrxn1α ^C/C^;Olig3^Cre/+^mutant mice do not h ave a deficit in updating or representing choice values (2-way RM ANOVA). (H-J) Nrxn1α ^C/C^;Olig3^Cre/+^ mutants exhibit a normal relationship between choice values and decision behavior. (K-M) Nrxn1α^C/C^;Olig3^Cre/+^ mutants do not exhibit a deficit in the allocation of choices guided by relative choice costs (K, 2-way RM ANOVA, left; two-sample t-test, right, p>0.05). Mutants exhibit no difference in task engagement (L, p>0.05) or in choice latencies (M, p>0.05). All data represented as mean ± SEM.

### Characterizing value-related neural signals within the dorsal striatum

Our data suggests that both constitutive Nrxn1α and forebrain excitatory neuron-specific mutants exhibit inefficient choice behavior driven by deficits in value updating. To explore whether these changes manifest at the level of neural activity within key circuits supporting reinforcement learning, we focused on direct pathway neurons of the dorsal striatum. We reasoned that this site is: (1) a common downstream target of cortical populations representing value and dopaminergic neurons conveying reward prediction errors (Bari et al., 2019; Bradfield et al., 2013; Parker et al., 2019; Parker et al., 2016); (2) implicated in action value coding (Kravitz et al., 2012; Samejima et al., 2005; Tai et al., 2012). To enrich for striatal dSPNs in our *in vivo* recordings of Ca^2+^ activity, we selectively expressing GCamp6f in projection inputs to the substantia nigra reticulata (SNr), via combined injection of retroAAV2.EF1α-3xFLAG-Cre in the SNr and AAV5.hSyn-DIO-GCamp6f in the dorsal striatum of control NEX^Cre^ mice (Fig.7A,B). Putative direct pathway SPNs (p-dSPNs) exhibited consistent Ca^2+^ activity patterns in relation to three task epochs that could be clearly assigned – trial start (center port light on), self-initiation (center port entrance) and choice/reward delivery (side port entry) (Fig.7C).

**Figure 7.**
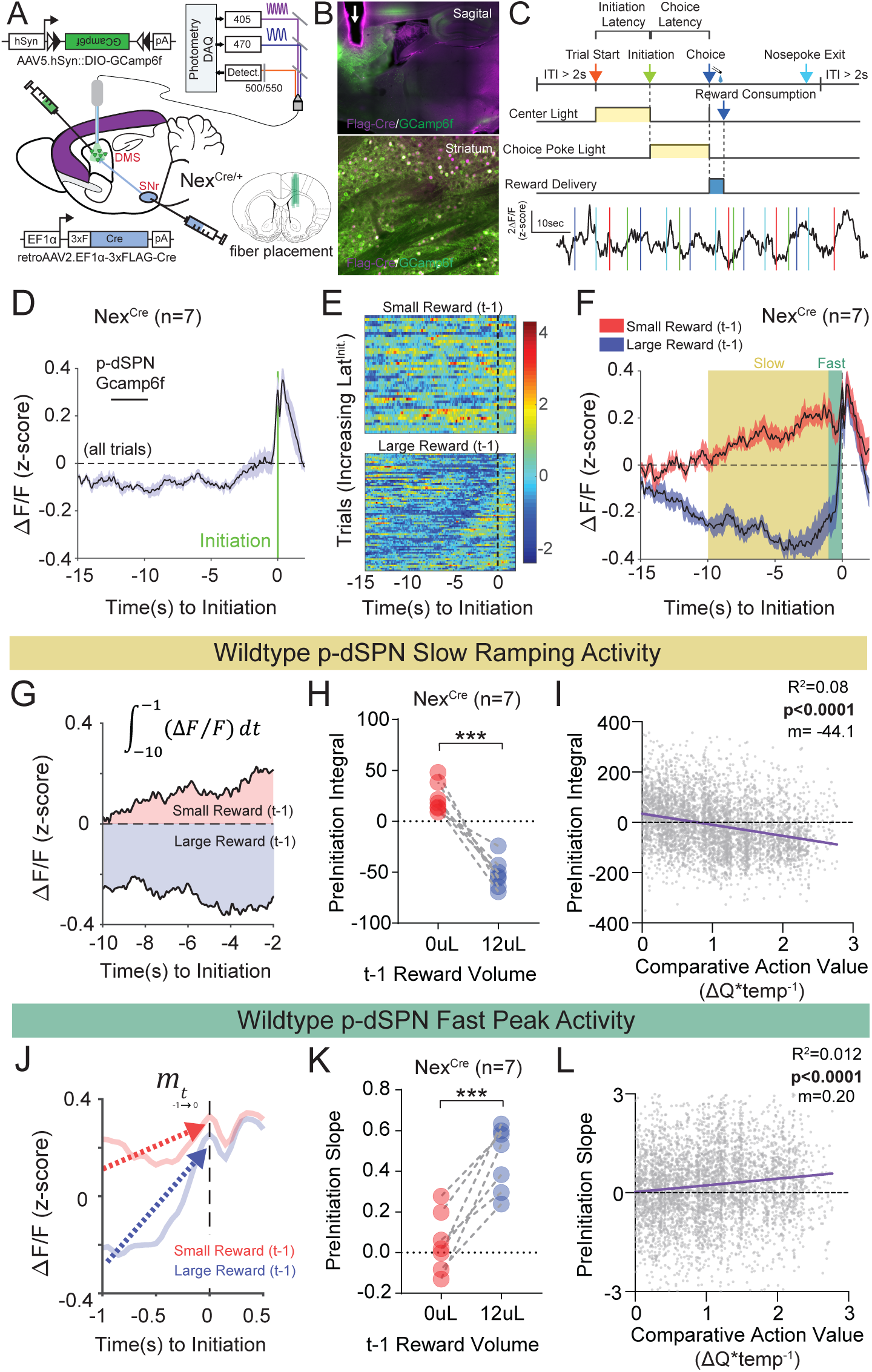
Quantifying value correlates in putative direct pathway SPNs of the dorsomedial striatum. (A) Schematic of experimental scheme. Control (Nrxn1α ^+/+^; Nex^Cre/+^, n=7) mice were injected with a retro-AAV2-EF1α-3xFLAG-Cre virus in the substantia nigra, pars reticulata (SNr). Ipsilateral injectio n of Cre-dependent GCamp6f allowed for enrichment of putative direct pathway SPNs (p-dSPNs). (B, top) Sagittal section of Nex^Cre^ brain showing GCamp6f expression in dorsal striatal SPNs and placement of 400µm optic fiber (white arrow). (B, bottom) Magnified view of striatum showing colocalization of nuclear FLAG-Cre and cytoplasmic GCamp6f. (B, bottom left) Location of fiber placements in Nex^Cre/+^. (C, top) Trial schematic and relationship of specific task epochs with p-dSPN Ca^2+^ signal (bottom). (D) Peristimulus time histogram (PSTH) of F/F for Nex^Cre/+^aligned to initiation event (all trials). The initiation of the action sequence (green bar) is associated with a rise in p-dSPNs activity. (E) Representative heat map of individual animal trials segregated by reward outcome on (t-1) trial (sorted by the latency to initiate). Trials following a large reward have greater signal suppression than those following small reward. (F) PSTH of F/F for Nex^Cre/+^ aligned to initiation event (segregated by outcome on (t-1)). Preinitiation p-dSPN dynamics exhibit two components – a slow ramping phase (yellow, *time*_-10→1_) followed by a fast spike phase (green, *time*_-1→init_), both of which are modulated by (t-1) reward outcome. (G) The slow → ramping phase is quantified by the integral of GCamp signal −10s to −1s before initiation. (H) There is a significant effect of (t-1) reward volume on the preinitiation integral during the slow ramping phase with large rewards showing greater silencing of p-dSPN activity (paired t-test, ***p=0.0002). (I) Preinitiation integral inversely correlates with the comparative action value of the upcoming trial, which is calculated using probability estimates from fitted reinforcement learning models and reflects the disparity in choice value on a trial to trial basis. (J) The dynamics of the fast peak phase are represented by the average slope of GCamp signal from −1s till initiation. (K) There is a significant effect of (t-1) reward volume on preinitiation slope during the fast peak phase (paired t-test,***p=0.0006) with large rewards showing steeper subsequent preinitiation slopes. (L) Preinitiation slope positively correlates with the comparitive action vlaue of the upcoming trial. Taken together, choice values regulate both fast peak and slow ramping activity in p-dSPNs.

Recent population Ca^2+^ imaging of striatal SPN populations has revealed a prolonged ramping activity prior to action sequence initiation (London et al., 2018). Together with our data (Fig.2) and other work documenting the modulation of initiation latency by prior outcome (Bari et al., 2019), we investigated the preinitiation window as one key timeframe for reward-related signals in dorsal striatal direct pathway neurons. An average of all trials aligned by initiation demonstrated multiple phases within the p-dSPN waveform (Fig. 7D). To understand how reward shapes p-dSPN activity, we segregated trials by previous (t-1) outcome. We found that most pre-initiation epochs following a ‘small reward’ trial had elevated activity compared to the population Ca^2+^ average, while trials following ‘large reward’ had suppressed activity relative to the population average (Fig.7E), a trend similarly present in the population data (Fig.7F).

To quantify signal dynamics, we took advantage of the bi-phasic nature of p-dSPN Ca2+ fluctuations, which exhibit both a slow ramping phase, occurring ∼10 seconds before an initiation, and a fast peaking phase, occurring 1-second before initiation. We found both signal components were differentially modulated by reward outcome: (1) for slow ramping, (t-1) large reward outcomes result in negative ramping or silencing of p-dSPN activity in comparison with small rewards (Fig.7G,H); (2) for fast peaking, larger rewards result in steeper peak activity as compared to smaller rewards (Fig. 7J,K). Furthermore, we noted significant correlations between both measures and trial-by-trial comparative action values (Fig.7I,L, see Methods), suggesting these p-dSPN signal components reflect components of value employed for future actions.

### Neurexin1α deletion in excitatory forebrain projection neurons disrupts reward-associated activity of striatal neurons

To examine whether deletion of Nrxn1α from forebrain projection neurons disrupted value-related neural signals within striatum, we performed population Ca^2+^ imaging of p-dSPNs in both Nrxn1α ^+/+^; Nex^Cre/+^ (Nex-Control) and Nrxn1α ^C/C^; Nex^Cre/+^ (Nex-Nrxn1α ^cKO^) mice during our serial reversal task. While we did not uncover a difference for the slow ramp signal component between genotypes (Fig.8A-C), we found that the slope of the fast peak was consistently lower in Nex-Nrxn1α ^cKO^ (Fig.8D,E). Furthermore, this deficit was specifically associated with failure to increase peak activity in response to large reward volumes (Fig.8F,G). Together, these data suggest that forebrain projection-specific Nrxn1α mutants do not have global disruptions of striatal circuit dynamics, but instead, a specific outcome-associated reduction in fast peak activity prior to trial initiation.

**Figure 8.**
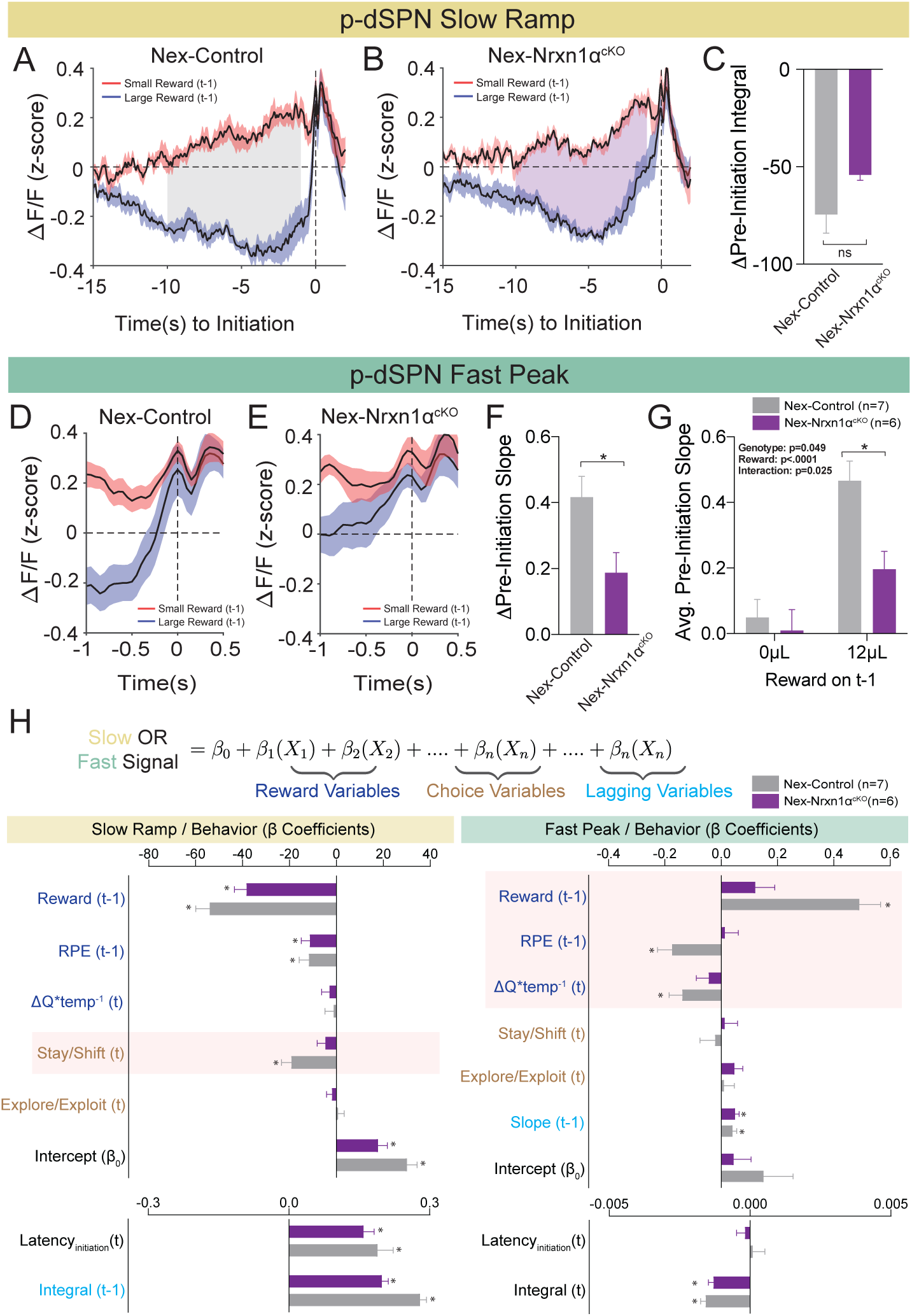
Restricted forebrain excitatory neuron deletion of Neurexin1α produces **a deficit in fast peak activity in p-dSPNs of the DMS**. (A,B) PSTH of F/F for Nex-control (Nrxn1α ^+/+^;Nex^Cre/+^, n=7, gray) and Nex-Nrxn1α ^cKO^ (Nrxn1α ^C/C^;Nex^Cre/+^, n=6, purple) mice, respectively, aligned to initiation ev ent (segreg ated by outcome on t-1). Shaded region corresponds to the difference in the preinitiation integral following large and small reward outcomes. (C) There is no statistically significant difference between Nex-control and Nex-Nrxn1α ^cKO^ in the integral of large versus small rewards (two-sample t-test, n.s., p=0.084). (D,E) PSTH of F/F for control and mutant animals, respectively, in the fast peak phase of preinitiation activity. (F) Nex-Nrxn1α ^cKO^exhibit smaller disparity in fast peak signals after unique reward outcomes, as evide nced by significant effect of genotype on slope of the fast peak (two-sample t-test, *p=0.025). (G) This difference in slope arises from a blunted GCamp response in mutants to large reward outcomes (2-way RM ANOVA). (H) Modeling Ca^2+^ signal dynamics as function of reward variables (blue), prior/future choice (gold), and lagging regressors (light blue), to capture prior circuit states. Value modulation of fast peak activity is blunted in Nex-Nrxn1α ^cKO^ mice (highlighted red box), while other components of the signal remain intact. Sl ow ramping is largely intact in mutant animals. All data represented as mean ± SEM.

To better understand whether mutation-associated changes in striatal neural signals related to specific components of the value-based decision-making process, we developed a linear-mixed effects model to explain variability in the integral of the slow ramp or the slope of the fast peak p-dSPN activity. Our model included variables accounting for reward processing (prior trial reward outcome and reward prediction error, disparity in action value between choices in the upcoming trial), choice behavior (choice, explore-exploit and stay-shift strategies), task engagement (initiation latencies) and lagging regressors to reflect “carry-over” plasticity effects from neighboring trials (Fig.8H, see STAR Methods). Surprisingly, we found that the blunting of the fast peak slope in Nex-Nrxn1α ^cKO^ mutants was specific to aspects of reward processing - i.e., while peak slopes had significant correlation to reward history, reward prediction error and comparative choice value in wildtype mice, these outcome-sensitive signal components are not present in mutant striatal population dynamics (Fig.8H). In contrast, reward-associated signal components are preserved in the mutants during slow ramping (Fig.8H), supporting a circumscribed alteration in striatal value coding. Together, these data demonstrate disrupted reward responsive activity in striatal circuits upon ablation of Nrxn1α in excitatory forebrain neurons. These activity changes align with our prior behavioral analysis showing Nrxn1α knockout in frontal projection neurons produces lower learning rate and sensitivity to outcome magnitudes (Fig.5E,F), generating smaller Q value discrepancies (Fig.5G).

## DISCUSSION

Understanding genetic contributions to brain disease requires bridging the sizeable chasm between molecular dysfunction and behavioral change. While behaviorally-circumscribed neural circuits provide a logical intermediary substrate, it has been challenging to identify disease-relevant neural populations owing to multiple factors: (1) difficulty in finding assays that provide stable, trait-like readouts of relevant behavioral constructs; (2) incomplete understanding of the specific computational algorithms and neural circuit implementations for behavioral constructs; (3) challenges localizing relevant neural circuits wherein gene perturbations drive behavioral dysfunction; (4) limitations in correlating mutation-associated patterns of neural activity with abnormal implementation of behavior.

Here we addressed these obstacles while investigating value-processing deficits in mice harboring mutations in Neurexin1α (Nrxn1α), a synaptic adhesion molecule associated with numerous neuropsychiatric disorders (Dachtler et al., 2015; Duong et al., 2012; Huang et al., 2017; Kirov et al., 2009; Rujescu et al., 2009; Sanders et al., 2015; Südhof, 2008). We found that constitutive Nrxn1α KO mice exhibited reduced choice bias towards larger benefits (modeled by greater reward volumes) and away from more costly outcomes (modeled by higher response schedules). Reinforcement modeling of choice behavior suggested altered mutant decision-making resulted from deficits in the updating and representation of choice value as opposed to how these values are transformed into action. Using a conditional allele and cell-type specific Cre recombinase drivers, we demonstrated that deletion of Nrxn1α from forebrain projection neurons, but not thalamic neurons, was able to recapitulate m ost aspects of the reward processing deficits observed in constitutive Nrxn1α KOs. Finally, we investigated how circuit-specific Nrxn1α mutants altered value-related neural signals within direct pathway neurons of the dorsal striatum. We found that while fast peak Ca^2+^activity immediately preceding trial initiation strongly reflected aspects of prior and current action value in wildtype mice, value-coding signals were disrupted in forebrain-specific Nrxn1α mutants.

### Deficits in Value-Based Action Selection in Neurexin1α Mutants

Reframing the study of disease-associated behaviors into endophenotypes is a powerful approach to revealing underlying genetic causality. Nevertheless, the study of disease-relevant cognitive endophenotypes in mice has proven challenging. Here we employed a feedback-based, two-alternative forced choice task that forces value comparisons between choices of differing reward magnitude. We believe this task has many advantages for investigating cognitive dysfunction associated with broad neuropsychiatric disease risk genes such as Nrxn1α produces stable within-mouse measures of benefit and cost sensitivity (Alabi et al., 2019), an ideal setting for observing between-group differences required in making genotypic comparisons. Second, it probes how outcome value is used to direct future action selection – a core neural process perturbed across many of the brain disorders in which Nrxn1α mutations have been implicated (Dichter et al., 2012; Gillan and Robbins, 2014; Maia a nd Frank, 2011).

We find that global deletion of Nrxn1α resulted in a persistent deficit in outcome-associated choice allocation, driven b oth by reductions in win-stay and lose-shift behavior (Fig.1C-E). Interestingly, similar reductions in win-stay behavior during feedback-based tasks have been demonstrated to drive choice inefficiency in both schizophrenia (Saperia et al., 2019) and autism (Solomon et al., 2015), disorders for which Nrxn1α has been consistently implicated. We observed that this value-related dysregulation manifests not only for the selection of higher-benefit actions, but also in the selection of less costly choices (Fig.3), as well as the outcome-dependent modulation of task engagement as read out by initiation latency (Fig.2). Together, these data converge to suggest Nrxn1α mutations either disrupt the normal function of brain circuits that internally represent value or the circuits that transform encoded values into actions.

### Deficits in the Updating and Representation of Value are Core Computational Deficits in Neurexin1α Mutants

In order to reveal which aspects of the decision process were altered in Nrxn1α mutants, we took advantage of prior work using Q-learning models to quantitatively describe relevant drivers of choice in feedback-based reinforcement paradigms (Daw, 2011; Sutton and Barto, 1998). Our data suggest that choice abnormalities in Nrxn1α KO mice reflect deficits in the updating of choice values, encapsulated by reductions in the learning rate (α) and outcome differentiation (γ) parameters, as opposed to differences in how mice translate value into action (β) or perseverate on actions m independent of outcome (κ) (Fig.4). These data are reminiscent of work from schizophrenic patients in a probabilistic reinforcement learning paradigm, where modeling suggested a reduction in the learning rate in patients versus neurotypical controls (Hernaus et al., 2018). Of particular interest, these investigators interpreted alterations in learning rate not to reflect perturbations in the reward prediction error (RPE) signal itself but to changes in how those signals were integrated to update value for future actions (Hernaus et al., 2019; Hernaus et al., 2018). While we are hesitant to directly map parameters of the reinforcement model to neural circuits, this interpretation suggests potentially relevant circuit loci might be those tasked with integrating dopaminergic RPE signals, including connectivity between cortical regions (representing internal/external state) and the input nucleus of the basal ganglia.

### Deletion of Neurexin1α from Forebrain Excitatory Neurons Recapitulates Choice Abnormalities of the Constitutive Knockout

The above hypothesis, together with robust expression of presynaptically-expressed Nrxn1α throughout cortex, directed us towards probing its function in these circuits. A large literature has implicated cortical regions in flexibly encoding the expected value of anticipated reward (Kennerley and Walton, 2011; Rolls, 2000; Tremblay and Schultz, 1999; Wallis and Kennerley, 2010; Wallis and Miller, 2003), reward-dependent modulation of working memory (Wallis and Kennerley, 2010) and forming associations between motivated behaviors and their outcomes (Hayden and Platt, 2010). Consistent with this, deletion of Nrxn1α from embryonic forebrain excitatory neuron progenitors recapitulated the value-based deficits observed in the constitutive KOs (Fig.5). While we do not claim this as the sole circuit-specific deletion capable of generating this phenotype, some degree of specificity was demonstrated by the striking absence of decision-making phenotypes in our thalamic Nrxn1α deletion (Fig.6).

Unfortunately, the broad recombinase expression of the Nex-Cre transgenic precludes us from assessing the importance of Nrxn1α in specific cortical areas. Co-expression networks seeded by autism candidate gene s have highlighted human mid-fetal deep layer cortical neurons from both prefrontal and primary motor/somatosensory cortices as potential sites of autism pathogenesis (Willsey et al., 2013). Furthermore, human patients with damage to the ventromedial prefrontal cortex exhibit similar deficits in value-based decision-making tasks as those seen in our Nex deletions (Camille et al., 2011; Fellows and Farah, 2007). Further assessment of the contribution of prefrontal Nrxn1α function to the observed phenotypes awaits Cre transgenic lines with both greater cortical regional specificity and embryonic expression.

### Circuit-Specific Ablation of Neurexin1α Abrogates Value-Associated Neural Signals within Striatum

Based on our behavioral data and computational modeling from multiple Nrxn1α mutants, well-documented expression patterns of Nrxn1α transcripts (Fuccillo et al., 1. 2015) and the known synaptic function of this molecule (Anderson et al., 2015; Aoto et al., 2013; Etherton et al., 2009; Missler et al., 2003), we hypothesized that the observed value-based abnormalities were the result of altered synaptic transmission at key sites for integration of RPEs into action value coding. Putative circuit loci included: (1) connections within value-encoding cortical areas; (2) connections from these cortical areas into striatum; (3) connections from cortex onto mesencephalic dopamine neurons that encode striatal-targeting RPE signals (Takahashi et al., 2011). To constrain the parameter space for initial recordings, we recorded population Ca^2+^ activity of putative dSPNs via fiber photometry, reasoning that all aforementioned possibilities could be read out by neural signals within SPNs of the striatum (Fig.7A-C). In support of this idea, we observed value-related signals leading up to trial initiation (Fig.7D,F; Fig.8A,D,H), consistent with population Ca^2+^ imaging signals observed in both SPN subtypes as mice approach palatable food (London et al., 2018). While our imaging does not provide the clarity of cellular-level approaches (Donahue et al., 2019; Kwak and Jung, 2019), it clearly demonstrates two phases of activity – a slow ramp occurring ∼10sec. before trial initiation and a fast peak in the 1sec. leading up to initiation – that correlate with prior reward outcome and RPE (Fig.7,8). Interestingly, the Nex-Nrxn1α mutants displayed a clear disruption of these reward variable correlations with p-dSPN activity, specifically for the fast peak immediately preceding trial initiation (Fig.8H). We suggest these data support a hypothesis wherein RPE signals are not appropriately integrated in Nex-Nrxn1α mutants, depriving striatal circuits of essential reward relevant information for subsequent action selection (Hernaus et al., 2019; Hernaus et al., 2018). Further *in vivo* neural recordings of corticostriatal circuits during this task together with input-specific interrogation of synaptic alterations are required to understand specifically how Nrxn1α mutations perturb communication of value-related information.

Extensive associations have been found between mutations in Nrxn1α and a range of neuropsychiatric disorders (Dachtler et al., 2015; Duong et al., 2012; Huang et al., 2017; Kirov et al., 2009; Rujescu et al., 2009; Sanders et al., 2015; Südhof, 2008). Here we show that Nrxn1α plays a key functional role in forebrain excitatory projection circuits that govern cognitive control of value-based action selection. It is interesting to speculate that the widespread nature of basic reinforcement learning abnormalities seen across neuropsychiatric diseases could be explained by similar network dysfunctions as those uncovered here for Nrxn1α mutants. Further work will be necessary to test the generalizability of these observations for other neurodevelopmental psychiatric disorders.

## ACKNOWLEDGMENTS

This work was supported by grants from the NIMH (F31-MH114528 to OA, R00-MH099243 and R01-MH115030 to MVF), the Whitehall Foundation and the IDDRC at the Children’s Hospital of Philadelphia. We thank Boris Heifets and Elizabeth Steinberg for assistance with initial Matlab code for photometry analysis. We also thank Alexandria Cowell for excellent assistance in mouse colony genotyping. Finally, we thank Patrick Rothwell and members of the Fuccillo lab for comments on the manuscript.

## AUTHOR CONTRIBUTIONS

Conceptualization, OA and MVF; Methodology, OA and MVF; Formal Analysis, OA; Investigation, OA, MF, MR; Writing – Original Draft, OA and MVF; Writing – Review and Editing, OA, MVF; Funding Acquisition, OA and MVF.

## STAR METHODS

### CONTACT FOR REAGENT AND RESOURCE SHARING

Further information and requests for resources should be directed to and will be fulfilled by the Lead Contact, Marc Fuccillo.

### EXPERIMENTAL MODEL AND SUBJECT DETAILS

Animal procedures were approved by the University of Pennsylvania Harbor Laboratory Animal Care and Use Committee and carried out in accordance with National Institutes of Health standards. Constitutive Neurexin1α (Nrxn1α) KO mice were obtained from the Südhof lab (Stanford University) (Geppert et al., 1998). Nrxn1α ^+/-^ males and females were bred to produce subject for this study. In sum, 11 Nrxn1α ^+/+^ and 12 Nrxn1α ^-/-^ mice were used in this study. 1 Nrxn1α ^-/-^ mouse died in the early stages of training and its results were excluded. Nrxn1α conditional knockout mice were generated from sperm stock (Nrxn1<tm1a(KOMP)Wtsi>) heterozygotes on the C57Bl/6N background) obtained from the MRC Mary Lyon Center (Harwell, UK). The lacZ gene was removed via crosses to a germline-FLP recombinase, which was then bred off, followed by at least 4 generations breeding to homozygosity within our colony. Nex^Cre^ mice (kind gift of Klaus-Armin Nave and Sandra Goebbels, Göttingen, Germany) were obtained and crossed onto Nrxn1α ^c/c^ mice (Goebbels et al., 2006). 11 Nex^+/-^ Nrxn1α ^-/-^, 14 Nex^-/-^ Nrxn1α ^c/c^ and 13 Nex^+/-^ Nrxn1α ^c/c^ mice were used in this study. Olig3^Cre^ mice were obtained (kind gift of Yasushi Nakagawa, University of Minnesota) and similarly crossed onto the Nrxn1α ^c/c^ colony (Vue et al., 2009). 8 Olig^+/+^; Nrxn1α ^C/C^ and 10 Olig^Cre/+^; Nrxn1α ^C/C^ mice were used in this study.

Whenever possible, animals were housed in cages with at least one littermate. One Neurexin1α wildtype and two Neurexin1α knockout animals were singly housed to avoid injury from infighting. Mice were food-restricted to maintain 85-90% of normal body weight and were given ad libitum access to water throughout the duration of the experiment. Mice were allotted 0.2-0.4 grams of extra food on non-experimental days to account for the discrepancy in caloric intake from not receiving reward in a task. A 7AM-7PM regular light-dark cycle was implemented for all mice used in this study. Cages were maintained in constant temperature and humidity conditions.

### BEHAVIORAL APPARATUS AND STRUCTURE

Experiments were conducted utilizing Bpod, a system specialized for precise measurements of mouse behavior (Sanworks LLC, Stony Brook, NY). A modular behavioral chamber (dimensions 7.5L×5.5W×5.13H inches, ID: 1030) with 3 ports capable of providing light cues and delivering liquid rewards was used to measure behavioral events. Each port was 3D printed from clear XT Copolyester and housed an infrared emitter and phototransistor to measure port entries and exits precisely. Behavior chambers were enclosed in larger sound-attenuating boxes. For each behavioral paradigm, illumination of the center port after a 1s intertrial interval indicated the beginning of a trial. Animals initiated trials by registering an entry to the lit center port, triggering a choice-period. The choice period was marked by the extinction of the center light and illumination of the ports on either side of the center. Mice were given an x-sec (varied by protocol) temporal window to enter either the left or right port and register a choice. Failure to register a choice in this period resulted in an omission, which was followed by a 3 second timeout and required the animal to reinitiate the task.

Successful registration of a choice resulted in the extinction of all port lights and the delivery of a variable volume of liquid supersac reward (3% glucose, 0.2% saccharin in filtered water) via a steel tube in the choice ports. Reward volumes and delivery probabilities were dependent on task conditions. The reward period lasted a minimum of 5 seconds. Following this mandatory minimum, the reward phase was extended if a mouse was noted to be occupying one of the three ports. The trial ended only after successfully confirmation of port exit from all three ports. Reward volumes were regulated via individually calibrated solenoid valves, with specific time/volume curves to deliver precise reinforcement.

All port entries, exits and other task events were recorded by the Bpod State Machine R1 (ID: 1027) and saved in MATLAB. Behavioral protocols and primary analysis were developed in MATLAB.

### OPERANT BEHAVIOR

#### Acquisition of Goal-Directed Contingency

Mice were habituated to behavior chambers and ports over a 3-day period. Each day, animals were given a 10-minute adjustment period followed by a program delivering 10µL of reward every 30 seconds for 40 min. The first 40 trials were grouped into 2 blocks, with reward delivered either from the left or the right port for 20 contiguous trials. Following this period, reward was alternated between left and right port for the remaining 20 trials. Port lights were illuminated for a 10 second period to indicate reward delivery, followed by a 20 second ITI.

Following this introductory period, mice were introduced to a goal-directed task that required them required them to acquire a light-chasing reward contingency. Trials were initiated as described previously. During the choice phase, one of the two lateralized ports was illuminated at random. Mice were given 10 seconds to register a choice, or an omission was charged. If entries into the unlit lateral port or the center port were registered a 3 second timeout occurred and the animals had to reinitiate the trial until they selected the correct port. Successful selection of the correct port resulted in 10uL of reward (P_rew_= 1.0). Sessions lasted 1 hour with no trial number limits. After 10 sessions, mice that had completed 2 consecutive days of >125 trials or 1 day >200 trials progressed to the serial reversal task. If mice missed this deadline, they were again assessed after their twelfth session. No mice that failed to meet these criteria by the twelfth session.

#### Serial Reversal Value Task

After successfully acquiring the action-outcome contingency described above, mice progressed to a forced-choice two-alternative serial reversal paradigm with variable reward outcomes. Trial initiation occurred as described above, via entry into the central port. To ensure accurate initiation latencies, the state of the center port was assessed after the ITI. The beginning of a trial was delayed if a mouse was found occupying this port. Initiation of a trial led to a 5sec choice period in which both left and right lateral ports were illuminated as choice alternatives. Following selection, a variable volume of reward was delivered contingent upon current task conditions (P_rew_= 0.75 and 0.4 were used here). The reward phase lasted 5sec and trial termination did not occur till after mice successfully disengaged from all ports. One Nrxn1α ^-/-^ mutant animal was excluded from the reversal study due to mis-calibrated solenoid valves.

Similar to our previous study, a “moving window” of proximal task events was used to monitor mouse choice patterns (Alabi et al., 2019). Changes of choice-outcome contingencies were initiated when 8 of the last 10 actions were allocated to the large reward volume side. Following detection of this event, the lateralization of reward volumes was switched. These contingency reversals were un-cued and served to mitigate outcome-insensitive behavior. Reward probabilities were the same for both choices and consistent over a given session. The relative reward contrast was consistent over a given session. Eight reward environments were tested (four relative reward ratios across two reward reinforcement rates). Animals performed the eight tests in a random sequence, performing the high reinforcement sessions before the low reinforcement sessions. For initial introduction to task structure, mice were run in the reversal paradigm (12µL v. 0µL) for 5-8 days prior to initiating the sequence of behaviors described above. All sessions were limited to 1 hour with no cap on trial number. Reward, however, was limited to 2000 µL in a session.

To ensure that behavioral measures were not overly influenced by spatial bias developed in one session (which could last for many subsequent sessions, across reward environments), sessions with excessive or carryover bias were excluded from this study and triggered a re-training phase before the experiment was continued. Bias was calculated as:

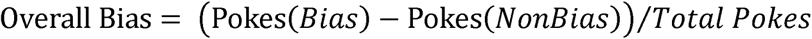

where Pokes (Bias) denotes the number of port entries to the side which received more pokes and Pokes (NonBias) represents the number of pokes to the side that received less. A bias exceeding 0.45 initiated an automatic re-training phase lasting at least one session. Sessions with biases > 0.2 triggered a watch-period in mice. If another session produced a bias >0.2 to the same spatial choice alternative, that session was marked as having carry-over bias from a previous session and excluded – also triggering a retraining phase. Sessions were additionally excluded if animals met 3 conditions in a single session: 1) overall bias exceeding 0.45; 2) failure to complete a minimum of 2 contingency switches; 3) failure to complete at least 100 selections of the nonbiased alternative. During re-training, animals performed one session of the 12µL v. 0µL reversal task to eliminate spatial bias.

#### Static Contingency Effort Task

A behavioral paradigm with a stable reward contingency over 150 trials was used to assess how costs shape behavior. Cost was modeled as increased operant responding (FR3) before delivery of a reward. Costs were applied to one alternative for 150 trials, following which a relative reward reversal was initiated (10µL v. 0µL) to eliminate the spatial bias developed during the task. Entry into one port during the choice phase led to extinction of the contralateral light. The chosen port remained lit until the animal completed the repetitive motor requirement necessary to obtain reward. Immediately upon completion of this requirement, reward was delivered as described previously. Equal reward volumes (8µL, P_rew_= 1) were implemented during the experimental phase of this task. Trial structure was the same as in the reversal paradigm described above. All sessions were limited to 1 hour. Each animal performed 2 experimental sessions to account for potential spatial biases. One with the high motor threshold on the right and the other with it on the left choice port.

Before animals were exposed to relative costs, they were acclimated to the new behavioral requirements by a three-session minimum training period in which they completed this task with an FR3 v. FR3 to increase response rate.

#### Cognitive Flexibility Assays

To measure cognitive flexibility, we employed an attentional set shifting task where the correct port was first indicated by a lit visual cue and subsequently switched to a fixed egocentric spatial position. Trials were structured as previously described. In the first 25 trials, a light cue denoted the position of reward. Mice initiated trials in which one of the lateralized alternatives was illuminated, at random, during a 10sec choice window. Selection of the illuminated port resulted in a 10µL reward, and selection of the unlit port resulted in a timeout. Following this baseline block, illumination of the choice ports continued to occur at random, but rewards were only delivered on one of the choice ports for the remainder of the session. Sessions were capped at 1hr and 250 trials.

To further probe behavioral flexibility, we utilized an egocentric spatial reversal task. Individual trial structure was preserved. In the first block of 25 trials, one of the choice ports was assigned as the reward port. Following this introductory block, the opposite port was assigned as the reward port. On each trial, one of the two ports was illuminated at random. A 10µL reward was given after selection of the appropriate port.

To account to potential biases and intersession fluctuations in performance, each animal was tested twice in each behavior – with alternating spatial cues in each session. P_rew_= 1 for both behaviors upon selection of correct alternative.

### ANALYSIS OF BEHAVIORAL PERFORMANCE

Data were analyzed using custom-written scripts developed in Matlab (Team, 2017). We utilized basic function supplemented by the following toolboxes: Bioinformatics, Curve Fitting, Data Acquisition, Global Optimization, Parallel Computing (Team, 2017). Analytical code is available on request.

#### Descriptive Parameters

The session performance index was calculated as:

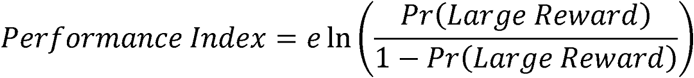

where *Pr(Large Reward)* refer to the percentage of total choice that animals made to the large reward alternative over the course of a session.

The relative reward-stay of an outcome, A, versus another outcome, B, was calculated as:

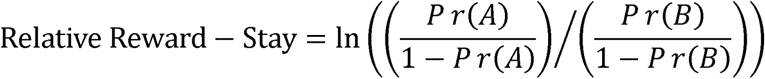

where Pr(A) and Pr(B) refer to the probability that mice stay on the choice alternative producing outcome A and B, respectively, on the *t-1* trial.

The adaptability index was calculated as:

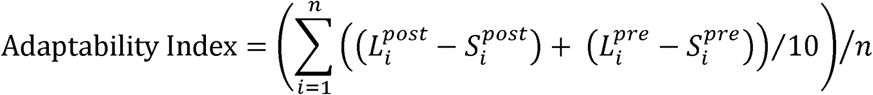

where 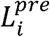 and 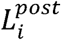 refer to the number of large alternative selections in the ten trials before and after the *i*th contingency switch in an individual session and 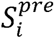 and 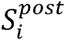 refer to the number of small alternative selections in the same time window. n is the number of blocks completed in a session.

The relative initiation latency was calculated as:

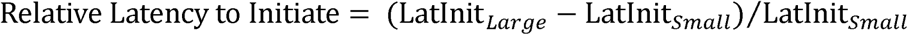

where LatInit_Large_ and LatInit_Small_ refer to the average latency to initiate trials following large reward and small reward outcomes, respectively, in an individual session.

#### Logistic Regression

We employed a logistic regression to model current choice as a function of past actions and outcomes (n=5 trials):

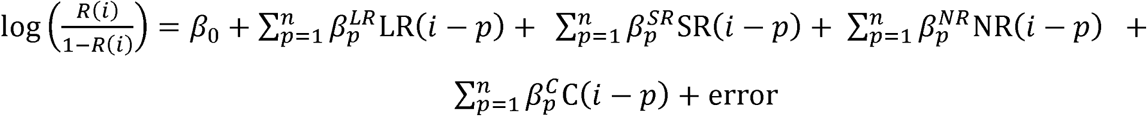

where *R(i)* is the probability of choosing the right-sided alternative on the *i*th trial. LR(*i-p*), SR(*i-p*) and NR(*i-p*) refer to the outcomes of the *p*th trial before the *i*th trial. LR(*i-p*) is defined such that LR(*i-p*) −1 if an animal received a large reward resulting from a right press on the *p*th previous trial, a −1 if an animal received a large reward resulting from a left press on the *p*th previous trial and 0 if the animal did not receive a large reward on that trial. SR(*i-p*) and NR(*i-p*) are defined similarly for trials that resulted in small reward and no reward outcomes, respectively. C(*i-p*) is an indicator variable representing the previous choice behavior of the mouse (C=1 for right-sided choice, C=0 for left-sided choice). These variables provide a complete accounting of the choice, reward history and interaction of the two in our task. This method assumes equivalent reinforcement from outcomes regardless of the lateralization of choice. The model was fit to 6 random blocks of 85% of choice data. The coefficient produced by these blocks were averaged to produce individual coefficients for each animal. Regression coefficients were fit to individual mouse data using the *glmfit* function in Matlab with the binomial error distribution family. Coefficient values for individual mice were averaged to generate the plots shown in the supplemental figures.

#### Reinforcement Learning Model

An adapted Q-Learning Reinforcement Model with 5 basic parameters was fit to the behavioral data produced by the relative reward serial reversal task (Daw, 2011; Sutton and Barto, 1998). Mouse choice patterns and outcome history were the primary inputs of the model. In order to capture trait-like characteristics of mouse behavior, behavioral sessions from the high and low reinforcement rate environments (4 sessions each) were grouped and entered into the model together. The values of the lateralized choice alternatives were initiated at 0 and updated as follows:

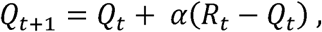

Where

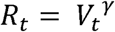

In this model, *Q_t_* is the value of the action taken on trial *t* and is the function that approximates the perceived reward volume resulting from that action. is defined as a compressive transformation of the reward volume, *V_t_*, delivered after a choice raised to the coefficient, γ. γ is the compression parameter that relates how sensitively mice respond to reward volumes of different magnitudes. *R_t_* – *Q_t_*, then, represents the reward prediction error (RPE) – the discrepancy between expected and realized reward – on trial *t*. The RPE is scaled by the learning rate (α), which determines the extent to which new information about the state-action pairing alters subsequent behavior. The scaled RPE is then used to update the value of the chosen action for the subsequent trial *t+1.* The value of the unchosen alternative was not altered on any trial and did not decay.

We utilized a modified softmax decision function to relate calculated action values with choice probabilities. The probability of choosing an alternative A on trial *t* was defined as:

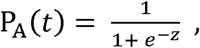

Where

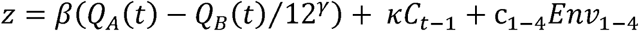

The inverse temperature parameter, β, is the conversion factor linking theoretical option values with realized choice output. High values β of indicate a tendency to exploit differences in action values, while lower values suggest more exploratory behavior. is the value of alternative A relative to the value of alternative B. In order to compare β across animals, this relative difference is scaled by 12^γ^, representing the maximum Q value (as largest delivered reward was 12µl). To account for the influence of proximal choice output on subsequent decisions, we included the parameter κ – the persistence factor. This measure captures the extent to which the animal’s choice on the *t-1* trial influences its choice on the *t* trial irrespective of outcome. is an indicator variable that denotes whether the animal selected alternative A on the previous trial (*C_t-1_* = 1) or if it selected alternative B (*C_t-1_* = −1). To account for potential differences in bias between sessions, a bias term, c_x_, with an indicator variable Env_x_, was added for each session that the animal perfomed. This constant term captures spatial biases that animals have or develop in the course of a behavioral session. We performed a maximum likelihood fit using function minimization routines of the negative log likelihood of models comprised of different combinations of our three parameters (α, β, γ, κ, c) in MATLAB (Vo et al., 2014). In order to resolve global minima, the model was initiated from 75 random initiation points in the parameter space.

### FIBER PHOTOMETRY

#### Viral Injection and Fiberoptic Cannula Implantation

Trained Nex^+/-^ Nrxn1α ^-/-^ (n=8) and Nex^+/-^ Nrxn1α ^c/c^ (n=6) mice were injected with adeno-associated viruses a nd implanted with a cus tom fiberoptic cannula on a stereotaxic frame (Kopf Instruments, Model 1900). Anesthesia was induced with 3% isoflurane + oxygen at 1L/min and maintained at 1.5-2% isoflurane + oxygen at 1L/min. The body temperature of mice was maintained at a constant 30°C by a closed loop homeothermic system responsive to acute changes in internal temperature measured via rectal probe (Harvard Apparatus, #50-722F). After mice were secured to the stereotaxic frame, the skull was exposed and anatomical landmarks bregma and lambda were identified. The skulls of the mice were subsequently leveled (i.e. bregma and lambda in the same horizontal plane) and 0.5mm holes were drilled on regions of the skulls above the target locations. A pulled glass injection need was used to inject 300nL of retroAAV2.EF1α-3xFLAG-Cre into the substantia nigra reticulata (SNr; AP: −4.2mm, ML: +/-1.25mm, DV : −3.11mm) followed by 500nL of AAV5.hSyn-DIO-GCamp6f into the dorsomedial striatum (DMS: AP: 0.85mm, ML: +/-1.35mm, DV: −2.85mm). Holes were drilled ipsilaterally and injections were performed unilaterally per mouse. Virus was infused at 125nL/min using a microinfusion pump (Harvard Apparatus, #70-3007) and injection needles were left in position for 10-20 minutes to allow diffusion of the viral bolus.

To implant each fiber optic, two 0.7mm bore holes were drilled ∼2mm from the DMS skull hole. 2 small screws were secured to the skull in these bore holes. A 400μm fiberoptic cannula was lowered into the DMS injection site. Small abrasions on the skull surface were created with a scalpel, following which, we applied dental cement (Den-Mat, Geristore A and B) to secure the fiber optic placement. After surgery, mice were given oxygen at 2L/min to aid in regaining consciousness. Mice were incubated for 4-6 weeks before recordings were performed. ∼2 weeks post-op mice were food deprived and reintroduced to the serial reversal task previously described.

#### Data Acquisition

Before recording sessions, mice were attached to a fiber-optic patch cord (400μm core, 0.48 NA; Doric Lenses) to enable recordings. Patch-cords were attached to a Doric 4-port minicube (FMC4, Doric Lenses) to regulate incoming and outgoing light from the brain. An LED light driver (Thor Labs, Model DC4104) delivered alternating blue (470nm, GCamp6f excitation) and violet (405nm, autofluorescence/movement artifact) light to the brain. Light was delivered at ∼50μW. The resulting excitation emissions were transferred through a dichroic mirror, a 500-550nm filter and were ultimately detected by a femotwatt silicon photoreceiver (Newport, Model 2151).

After attachment to the fiber-optic, animals were given a 5-min window to recover from handling before the initiation of a session. All recorded mice were trained to perform the relative reward serial reversal task before surgery. Animals were reintroduced to the task ∼2 weeks post-surgery. At 3 weeks, expression of the GCamp6f construct was assessed and animals were trained to perform the task with the attached fiber-optic. After a minimum of 4 weeks and 3 full training sessions with the fiber optic, animals were eligible for recordings. Sessions lasted 1 hour. We introduced a 0-1 temporal jitter after the ITI and before the choice period to aid in dissociating task events.

#### Signal Processing and Analysis

Raw analog signals from behaving mice were demodulated (Tucker Davis Technologies, RZ5 processor) and recorded (Tucker Davis Technologies, Synapse). Demodulated 470nm and 405nm signals were processed and analyzed using custom Matlab (MathWorks, R2018b) scripts that are freely available on request. Signal streams were digitally filtered and down-sampled to 20Hz. To account for de-bleaching of backround autofluorescence in the patch cords over long recording sessions, the demodulated 470nm and 405nm signals were fitted with a cubic polynomial curve, which was subsequently subtracted from the signal. The ΔF/F of the debleached signals were calculated and the 405nm control signal was subtracted from the 470nm GCamp6f emission signal. The subtracted ΔF/F was transformed into z-scores by subtracting the mean and dividing by the standard deviation of a 1min window centered on each point. These standardized fluorescence signals were used for all subsequent analysis and visualization. The Bpod State Machine delivered electronic TTLs marking behavioral events to Synapse Software, which recorded their time and direction.

#### Modeling Signal Dynamics

The dynamics of preinitiation signal components was modeled as function of action output in the form of upcoming choice behavior (choice lateralization relative to implant [*Choice*], stay/shift behavior [*Stay*], explore/exploit behavior[*Explore*]), reward (reward volume on previous trial [*RewardHist*], reward prediction error [*RPE]* on previous trial and the relative action value on the current trial [*Q*temp^-1^*]), prior signal dynamics (the preinitiation slope and integral on the previous trial [*PIS* and *PIT*, respectively,) and the latency to initiate trials [*LatInit*]. Because the slope occurs after the integral on every trial and because slope and integral components are anti-correlated, the preinitiation integral on the *t* trial was included as a regressor in the modeling of the slope component. To account for individual animal differences in preinitiation signal components, we utilized a linear mixed-model:

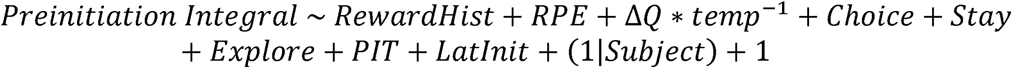

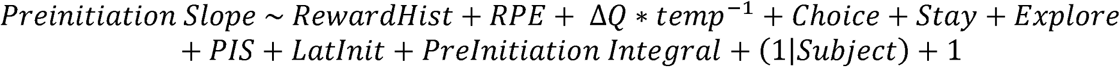

#### Histology and Immunohistochemistry

Mice were perfused via the left ventricle of the heart with 10mL of 90% formalin. Whole brains were isolated and post-fixed in formalin overnight. 50µm coronal and sagittal slices were sectioned in PBS. Slices from mice included in behavioral experiments were immediately mounted on microscope slides for imaging on an automated fluorescence microscope (Olympus BX63) at 10x (Olympus, 0.4NA). Additional sections were blocked in 3% normal goat serum in PBS for 1 hr and incubated with primary antibody overnight (1:500 Chick anti-GFP, abcam 13970; 1:1000 Mouse anti-FLAG, Sigma F1804). The following day, slices were washed with PBS and incubated for 3 hours with secondary antibody (1:1000 Goat Alexa488-conjugated anti-Chick, abcam 150173; 1: 1000 Goat Alexa647-conjugated anti-Mouse, Invitrogen #A-21235). Slices were washed 3x with PBS for 30 minutes and mounted on slides. Images were acquired from the same epi-fluorescent microscope as other images.

### STATISTICAL METHODOLOGY

All data were initially tested with appropriate repeated measure ANOVA (Prism8.0). Univariate regressions were performed in Prism8.0. Multivariate linear regressions were performed using the *fitlm* function in MATLAB. Multivariate linear mixed models were performed using the *fitlme* function in MATLAB. Main effect and interaction terms are described within figures, figure legends and the results. Preinitiation slope coefficients were calculated using the *polyfit* function in MATLAB. The integral of photometry signals was calculated using the *trapz* function in MATLAB.

