## Supplemental Figures for "Disruption of Nrxn1α within excitatory forebrain circuits drives value-based dysfunction"

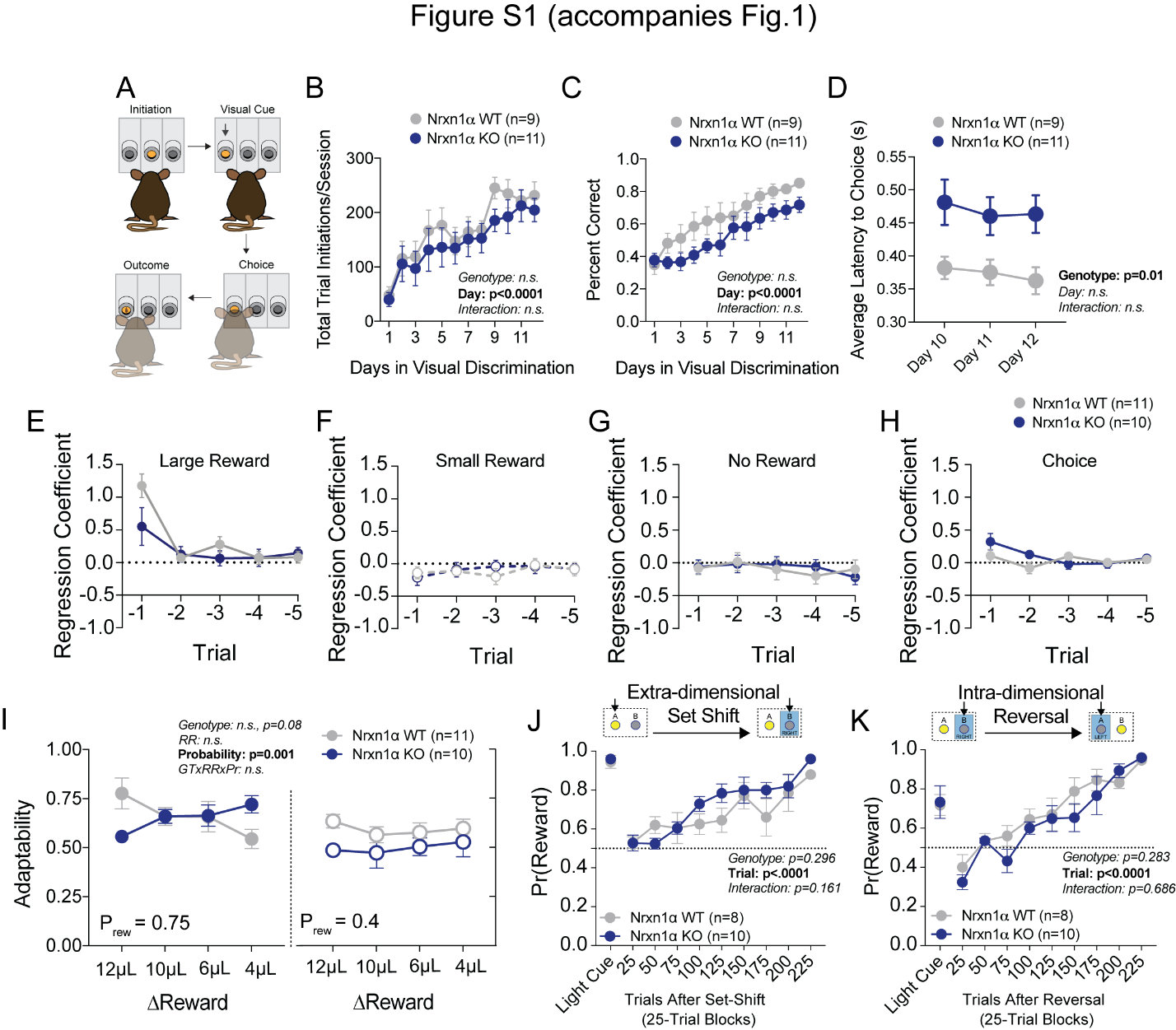

**Supplemental Figure 1.**

(A) Schematic of visual discrimination task. Mice acquired a simple goal-direction contingency over repeated sessions. (B) Task engagement was measured as the total number of registered trial initiations. Nrxn1α KO mice exhibit no difference in task engagement from wildtype littermates during acquisition of visual discrimination. (C) Performance was measured as the proportion of trial initiations that resulted in the selection of the lit port. Nrxn1α mice exhibit no deficit in visual discrimination as compared to wildtype littermates. (D) Nrxn1α KO animals exhibit extended choice latencies throughout the last 3 days of task acquisition. (E-H) Logistic regression coefficients for Nrxn1α wildtype and knockout mice (ΔReward = 12μL). The influence of past choice and reward outcome is heavily discounted after the t-1 trial. (I) There is a trend towards lower adaptability measures in Nrxn1α knockout mice in the relative reward reversal paradigm (see Fig.1A). (J) Nrxn1α KO animals exhibit similar performance to wildtype controls in extra-dimensional set shift where reward target switches from visual cue to ego-centric spatial cue. (K) Nrxn1α KO animals exhibit similar performance to wildtype controls in egocentric spatial reversal task. (B-D, I-K analyzed by 2-Way RM ANOVA). All data represented as mean ± SEM.

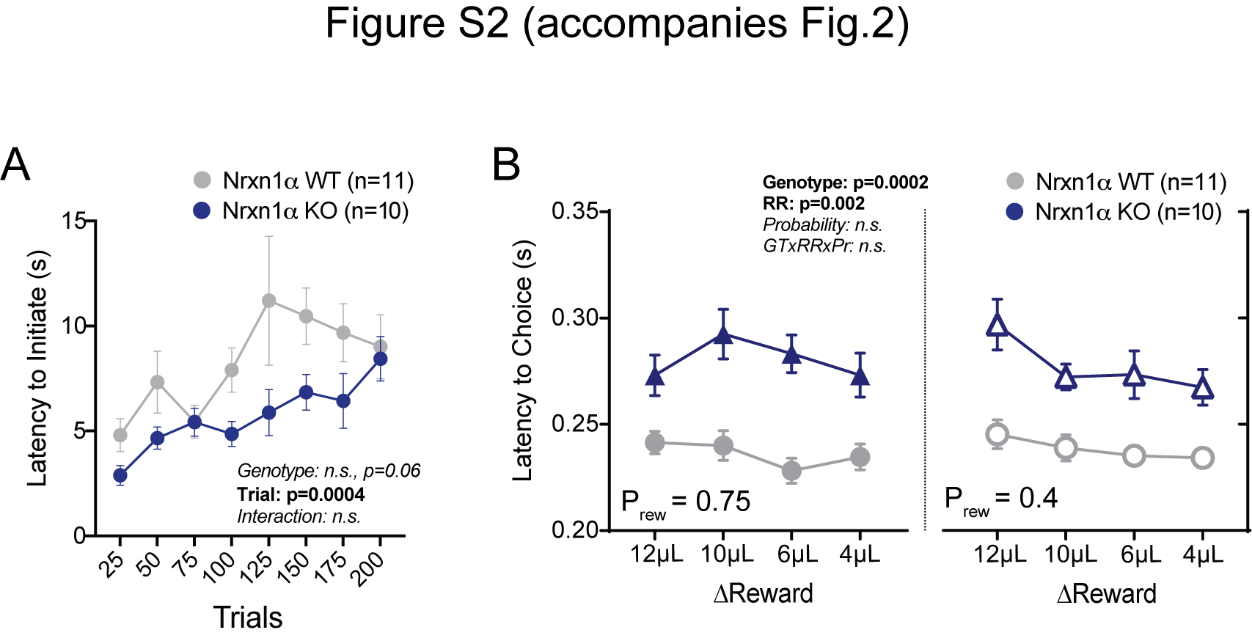

**Supplemental Figure 2.**

(A) The latency to initiate of animals increases as mice consume more reward in this task (ΔReward = 12μL). There is a trend towards lower initiation latencies in Nrxn1α KO animals. (B) Nrxn1α KO mice exhibit extended choice latencies across reward environments. All data analyzed by 2-Way RM ANOVA. All data represented as mean ± SEM.

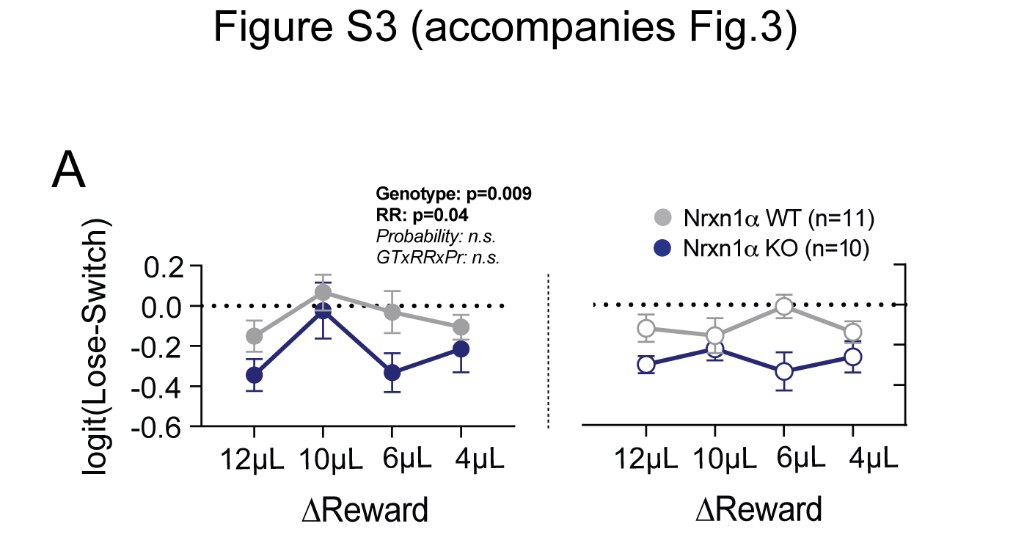

**Supplemental Figure 3.**

(A) Lose Switch. Nrxn1α mutant mice (n=10) exhibit a lower propensity to shift choice behavior after receiving no reward than wildtype littermates (n=11). Analyzed by 3-Way RM ANOVA. All data represented as mean ± SEM.

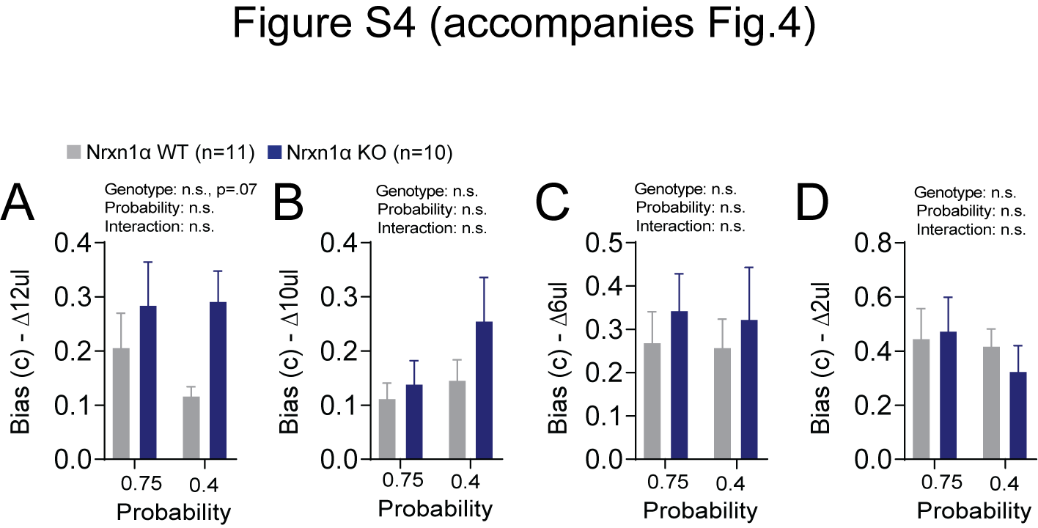

**Supplemental Figure 4.**

(A-D) Model Biases. A bias term was generated for each relative reward ratio to capture potential difference in how animals develop biases in different reward environments. There is no statistically significant effect for genotype or probability on the degree of bias generated (ΔReward = 12,10,6,4μL for A-D, respectively). All data analyzed by 2-Way RM ANOVA. All data represented as mean ± SEM.

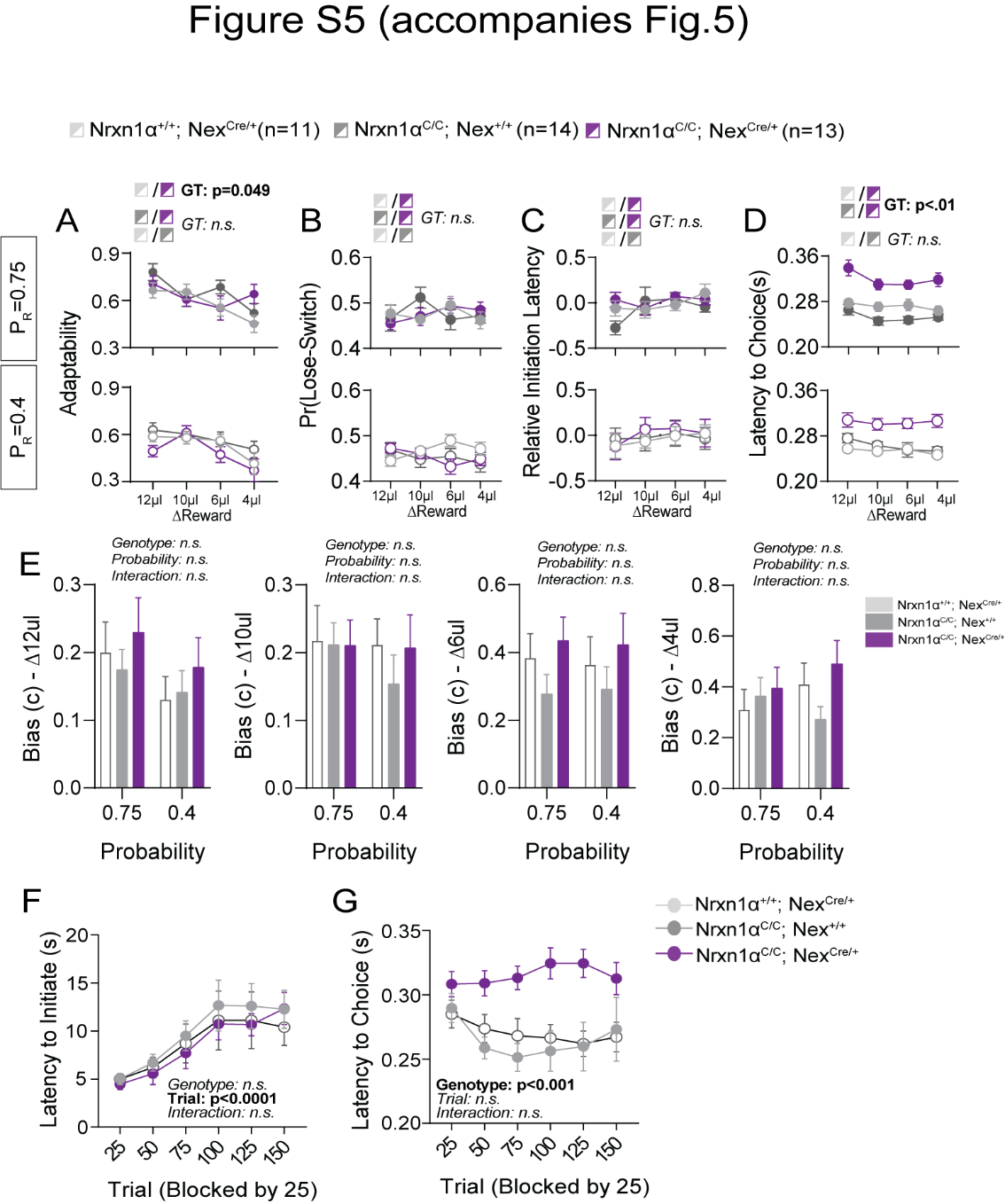

**Supplemental Figure 5.**

(A) In-task adaptability. There is a main effect for genotype on the adaptability measures between Nrxn1α^C/C^; Nex^Cre^ controls and mutant mice. Otherwise there are no differences in adaptability noted for remaining genotypic comparisons. (B) Lose-Switch. There is no effect for genotype on the probability that animals will alter their decision after receiving no reward. (C) Relative initiation latency. We noted no significant differences in the relative latency to initiate trials after distinct reward outcomes. We did not view the same robust modulation of task vigor by previous reward outcome in the Nrxn1α^C/C^ colony, precluding this line of analysis. (D) Choice Latency. Nex^Cre^;Nrxn1α^C/C^ mutant mice (n=13) exhibit longer choice latencies in comparison with Nex^Cre^ (n=11) and Nrxn1α^C/C^ (n=14) controls across reward environments. (E) Model Biases. There were no statistically significant effects of genotype or probability on the degree of bias generated (ΔReward = 12,10,6,4μL from left to right). (F) Initiation latency during choice paradigm. There is no statistically significant difference in the initiation latencies of mutants as sessions progress. (G) Nex^Cre^;Nrxn1α^C/C^ mutant animals exhibit extended choice latencies in comparison with their wildtype counterparts. (A-D analyzed by 3-Way RM ANOVA, E-G analyzed by 2-Way RM ANOVA).

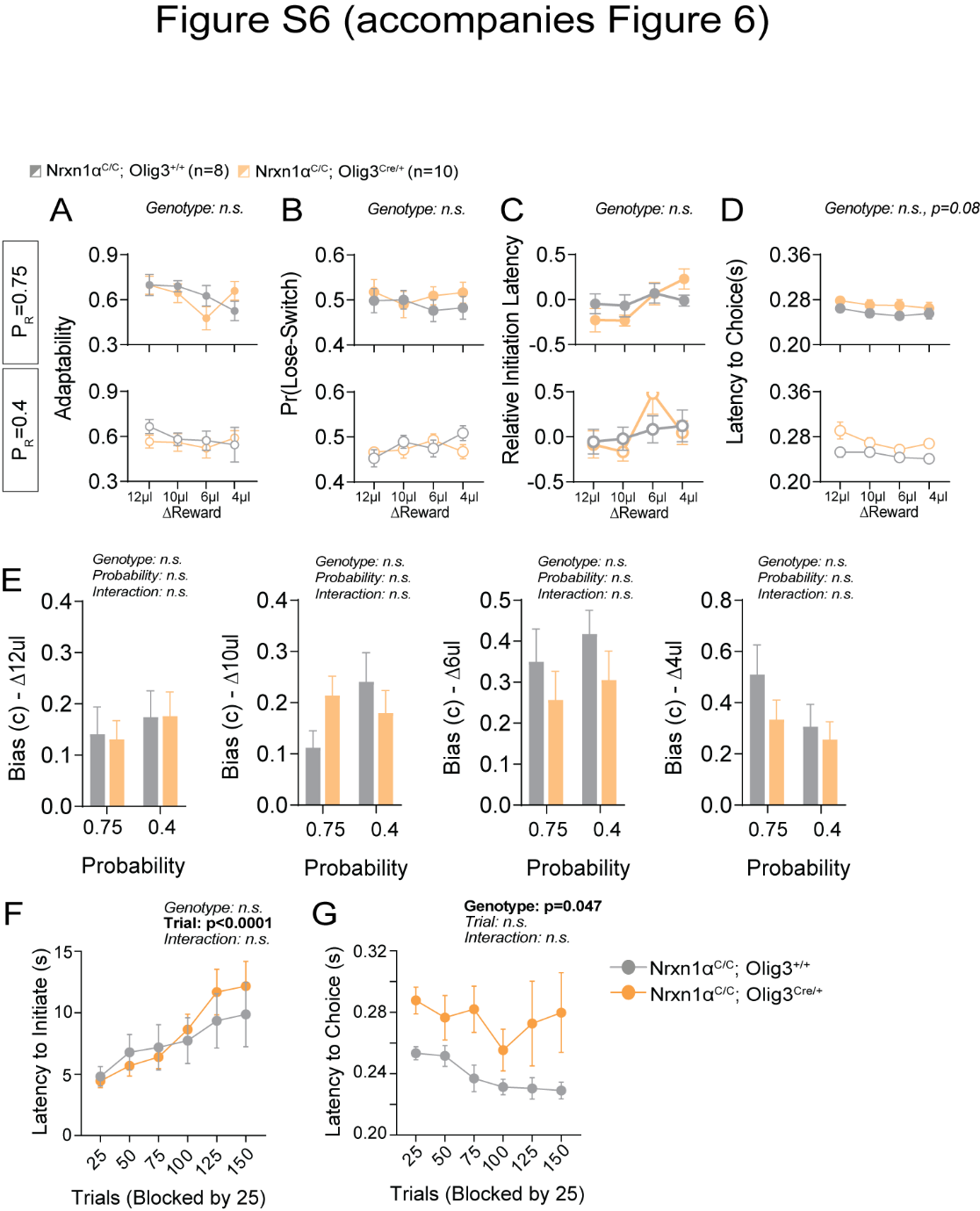

**Supplemental Figure 6.**

(A) In-task adaptability. There is no effect for genotype on the adaptability measures of Olig^Cre^ control and Olig^Cre^;Nrxn1α^C/C^ mutant mice. (B) Lose-Switch. There is no effect for genotype on the probability that animals will alter their decision after receiving no reward. (C) Relative initiation latency. We noted no significant differences in the relative latency to initiate trials after distinct reward outcomes. (D) Choice Latency. Olig^Cre^;Nrxn1α^C/C^ mutant mice (n=10) exhibit a trend towards extended choice latencies in comparison with Nrxn1α^C/C^ (n=8) controls across reward environments. (E) Model Biases. There is no statistically significant effect for genotype or probability on the degree of bias generated (ΔReward = 12,10,6,4μL from left to right). (F) Initiation latency during choice paradigm. There is no statistically significant difference in the initiation latencies between genotypes as sessions progress. (G) Olig^Cre^;Nrxn1α^C/C^ mutant animals exhibit extended choice latencies in comparison with their two wildtype counterparts. (A-D analyzed by 3-Way RM ANOVA, E-G analyzed by 2-Way RM ANOVA).
